# Genomic stop codon scanning reveals quantitative principles of nonsense-mediated mRNA decay

**DOI:** 10.64898/2025.12.20.695734

**Authors:** Michael A. Cortázar, Jacob Schmidt, Iman Egab, Zeynep Coban-Akdemir, Sujatha Jagannathan

## Abstract

Nonsense-mediated mRNA decay (NMD) degrades transcripts containing premature termination codons (PTCs), critically shaping the disease outcomes of protein-truncating variants. While existing NMD rules categorize PTCs as NMD-triggering or NMD-evading, they cannot quantitatively predict the degree of NMD activity for a given endogenous PTC variant. To provide quantitative insight into NMD, we used saturation genome editing (SGE) to systematically introduce all possible PTCs (TAA, TAG, TGA) at every codon position spanning the first, penultimate, and three internal exons of the Lamin A/C (*LMNA*) gene. Combining targeted sequencing with NMD inhibition, we measured mRNA expression and NMD activity for 722 PTCs and 211 single nucleotide variants (SNVs). Our data validate known positional trends in NMD activity but reveal unexpected complexity. In the penultimate exon, the PTC position effect extends beyond the binary 50-55 nucleotides (nt) rule, revealing a quantitative relationship between the PTC-EJC distance and NMD activity. At the 5’ end, NMD is completely absent in the first 21 codons of *LMNA*, followed by sharp activation over the next 4 codons. Both patterns, at the 5’ end and in the penultimate exon, are unexplained by current models. Finally, internal exons show robust NMD with outliers consistently mapping to predicted readthrough-permissive sequence contexts, including the conserved readthrough promoting TGA-CT motif. This comprehensive dataset provides an unprecedented resource for understanding the quantitative impact of PTC position and sequence context on NMD, with direct implications for the clinical interpretation of nonsense variants in the human population.

## Introduction

Mutations that introduce premature termination codons (PTCs) are responsible for approximately 20% of all disease-associated single-base pair substitutions (Mort et al., 2008). PTC-containing mRNAs are recognized and degraded by nonsense-mediated RNA decay (NMD) which prevents the synthesis of potentially toxic, truncated proteins (Campbell et al., 2023; Carrard & Lejeune, 2023; Kurosaki & Maquat, 2016; Udy & Bradley, 2022). However, because truncated proteins can retain residual function, NMD can also contribute to disease severity by eliminating otherwise partially functional transcripts. As such, NMD serves as an important modifier of many genetic disorders (Khajavi et al., 2006; Lindeboom et al., 2019).

According to the canonical model in mammalian cells, the NMD pathway primarily distinguishes between PTCs and normal stop codons through the existence of exon junction complexes (EJCs) downstream of the terminating ribosome (Kashima et al., 2006; Nagy & Maquat, 1998). During splicing, the exon junction complex (EJC) is loaded onto the mRNA 20–24 nt upstream of the exon-exon junction (Le Hir et al., 2000; Sauliere et al., 2012; Singh et al., 2012) and removed by the translating ribosome (Dostie & Dreyfuss, 2002; Lejeune et al., 2002; Sato & Maquat, 2009). When a ribosome prematurely stops at the PTC, communication with a downstream EJC through the NMD factor UPF1 and the translation termination factors eRF1/3 (Kashima et al., 2006) is thought to trigger mRNA degradation (Eberle et al., 2009; Gatfield & Izaurralde, 2004; Huntzinger et al., 2008; Loh et al., 2013; Unterholzner & Izaurralde, 2004).

Based on this model, PTCs in the last exon or within the last 50 nt of the penultimate exon do not trigger NMD, since the ribosome in this region is predicted to evict the last EJC before it stops at the PTC, termed the “50-nt rule” (Nagy & Maquat, 1998). In clinical genetics, the 50-nt rule is routinely used as a binary classifier to label PTCs as NMD-triggering or NMD-evading (Singer-Berk et al., 2023).

While the 50-nt rule provides a useful framework to interpret nonsense variants, it is not deterministic: at least 30% of PTCs that are predicted to be NMD-triggering based on this rule evade NMD (Lindeboom et al., 2016). Moreover, several genes show clear exceptions to the 50-nt rule, with poor mRNA degradation at PTC locations predicted to trigger NMD, including *HNF1B* (Harries et al., 2005), *HBB* (Danckwardt et al., 2002; Inacio et al., 2004; Neu-Yilik et al., 2011; Peixeiro et al., 2012), *HBA1* (Pereira et al., 2015; Silva et al., 2006), *TPI1* (Zhang & Maquat, 1997), *ATRX* (Howard et al., 2004), *MSH3*, *TAF1B*, and *TGFBR2* (You et al., 2007). NMD variability in reporters can be explained, at least in part, by an influence of the number of EJCs upstream and downstream of the PTC, or even exon length (Hoek et al., 2019).

Therefore, PTCs located upstream of the 50-nt boundary can be associated with a wide range of NMD activities. However, the current understanding of the NMD pathway does not permit reliable prediction of the magnitude of NMD for a given PTC based solely on its position within the mRNA or its local sequence context. It is also not known if the 50-nt mark reflects a sharp transition from strong to no NMD or how far upstream the influence of the last EJC on a PTC extends to enable efficient NMD.

A particularly striking exception to the 50-nt rule occurs at the opposite end of the open reading frame (ORF): PTCs near the translation initiation site (start-proximal PTCs) often evade NMD, despite being far upstream of the last exon-exon junction in a context where the 50-nt rule predicts efficient degradation. In a β-globin reporter mRNA, PTCs within the first 23 codons do not trigger mRNA depletion (Inacio et al., 2004; Neu-Yilik et al., 2011). This resistance has been attributed to translation reinitiation downstream of the PTC (Neu-Yilik et al., 2011), or stimulation of translation termination at the PTC by the cytoplasmic poly(A)-binding protein 1 (PABPC1) in a potential closed loop conformation of the mRNA (Peixeiro et al., 2012; Silva et al., 2008). The latter mechanism aligns with the faux 3’-UTR model (Amrani et al., 2004) in yeast, where EJC-independent NMD is triggered when translation termination occurs far from the poly(A) site or the normal termination region.

Factors affecting the kinetics of translation termination at PTCs are also thought to impact NMD. In yeast extracts, ribosomes appear to reside for a longer period of time at PTCs when compared to normal stop codons, suggesting slow or inefficient termination (Amrani et al., 2004), although extracts from HeLa cells did not show this effect (Karousis et al., 2020).

Consistent with this view, slow peptidyl tRNA hydrolysis, especially at glycine codons immediately upstream of a PTC, creates an extended translation termination window that enhances mammalian NMD activity (Kolakada et al., 2025). A C nucleotide immediately downstream of the PTC at the +4 position was also associated with differential NMD activity (Mabin et al., 2025). The three stop codons differ intrinsically in readthrough permissiveness (TGA>TAG>TAA), and the surrounding nucleotide context, particularly a C at the +4 position further enhances readthrough (Buckley et al., 2024; Manuvakhova et al., 2000; Wangen & Green, 2020). The efficient readthrough TGA-CT motif is a conserved element for the functional C-terminal extension of proteins across diverse species (Loughran et al., 2014; Stiebler et al., 2014). Yet, how these sequence features interact at endogenous PTCs to determine NMD activity has not been systematically studied.

Studies using human population RNA-seq data sets have validated the general applicability of the 50-nt rule and revealed a global trend of poor mRNA clearance for start-proximal PTCs (∼200 nt downstream of the translation initiation site, 200-nt rule) and long exons (>400 nt) (Lindeboom et al., 2016; Rivas, Pirinen, Conrad, Lek, Tsang, Karczewski, Maller, Kukurba, DeLuca, Fromer, Ferreira, Smith, Zhang, Zhao, Banks, Poplin, Ruderfer, Purcell, Tukiainen, Minikel, Stenson, Cooper, Huang, Sullivan, Nedzel, Consortium, et al., 2015; Teran et al., 2021). However, these studies rely on meta-analyses of aggregated data from nonsense mutations across different genes and cellular contexts, precluding systematic interrogation of NMD behavior along a single mRNA. Moreover, NMD activity is inferred indirectly: reduced transcript abundance is assumed to reflect degradation by NMD, overlooking the possibility that PTCs may also impair transcription (Stalder & Muhlemann, 2007) or mRNA processing (Abrahams et al., 2021). A direct and systematic quantification of NMD activity along the mRNA is therefore needed.

Here, we provide a quantitative map of mRNA expression changes and NMD activity along the LMNA mRNA transcript following systematic codon-by-codon CRISPR/Cas9 integration of PTCs. The *LMNA* gene is well-suited for this analysis: it is expressed and non-essential in HAP1 cells, spans 12 exons, allowing interrogation of the first, internal and penultimate exons, and is clinically relevant. Mutations in *LMNA* cause a broad spectrum of diseases, including cardiac, muscular and neurological disorders. Our results yield an unprecedented quantitative map of NMD strength and variability across the LMNA mRNA, demonstrating that PTC position is the major determinant of NMD efficiency. First, NMD is largely absent at both ends of the ORF: completely inactive in the first 21 codons with sharp activation over codons 22-25 and, in the penultimate exon, changing as a function of the distance between the PTC and last exon-exon junction. Second, NMD is robust and stable across internal exons, with variability limited to discrete drops at specific positions. Third, these outliers map to sequence contexts predicted to promote translation readthrough, implicating translation termination efficiency as a key modulator of NMD.

## Results

### A saturation genome editing (SGE)-based approach to systematically integrate all possible PTCs in *LMNA* exons

We applied SGE to systematically introduce all possible PTCs (TAA, TAG, TGA) at each codon across selected exons of *LMNA*. SGE enables pooled editing by co-transfection of the CRISPR-Cas9 system and a donor plasmid library, followed by homology-directed DNA repair (HDR) (Findlay et al., 2014; Findlay et al., 2018) (Fig. 1A). To test established NMD rules, we targeted exons spanning key regulatory contexts: exon 1 (start-proximal), exons 4, 7, and 10 (internal), and exon 11 (penultimate exon containing the 50-nt boundary) (Fig. 1B). Given the length of exons 7 and 11, we prioritized editing the 3’ end region to assess the potential influence of the nearest EJC. As controls, we included ClinVar-reported SNVs (mostly missense, some synonymous) that should not activate NMD. Each library also contained a fixed synonymous substitution at the Cas9 target site to prevent re-cutting after successful HDR (Findlay et al., 2014). We compared each PTC and SNV variant to an integrated synonymous substitution-only sequence context. HAP1 cells were edited with SGE, and, at day 12, treated with the small molecule NMD inhibitor, SMG1i (1μM for 4 hours) (Gopalsamy et al., 2012), or DMSO. Variant mRNA expression and NMD efficiency were quantified by targeted sequencing of the edited exon from both genomic DNA (gDNA) and RNA (Fig. 1A).

**Figure 1.**
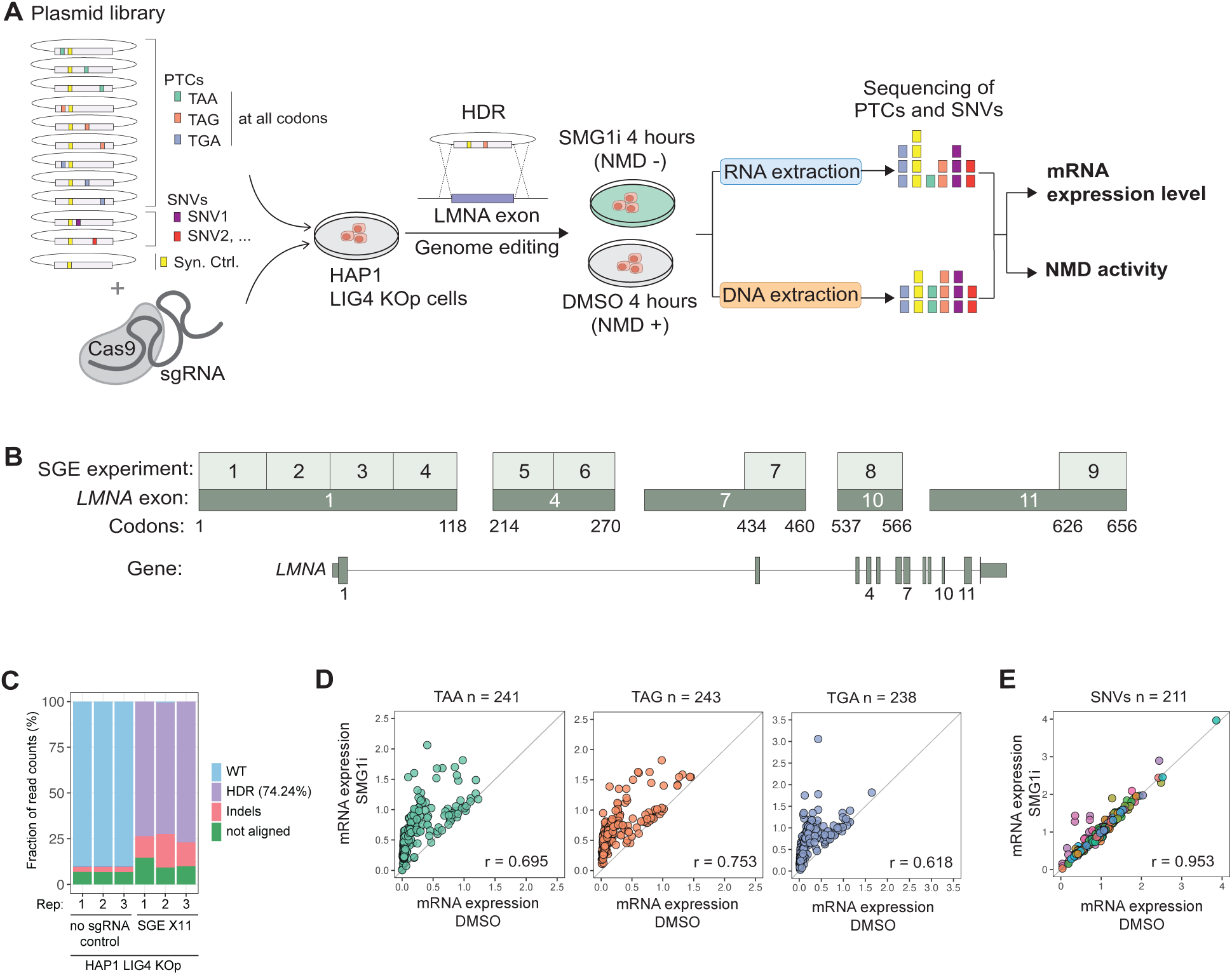
A saturation genome editing (SGE)-based approach to systematically integrate all possible PTCs in *LMNA* exons. **(A)** HAP1 cells are co-transfected with CRISPR/Cas9-encoding and donor HDR plasmids, followed by inhibition of NMD 12 days post-transfection and targeted sequencing of the SGE-treated region at both the DNA and RNA levels. **(B)** SGE target regions spanning exons across the *LMNA* gene. **(C)** Proportion of sequenced read tags from genomic DNA mapping to wild-type (WT; unedited cells), homology-directed repair (HDR), or insertions and deletions (indels), from the SGE experiment targeting *LMNA* exon 11 (X11) **(D, E)** Correlation of mRNA expression levels between DMSO and SMG1i conditions for each stop codon type (TAA, TAG, TGA) and SNVs.

To enrich low-frequency HDR events over non-homologous end joining (NHEJ), the ligase IV gene (*LIG4*) is routinely knocked out in SGE experiments (Findlay et al., 2018; Radford et al., 2023; Waters et al., 2024). However, because NMD can vary substantially across individual cells within a population, using a single-cell–derived clonal knockout could bias results. To avoid this limitation, we developed SelectRepair knockout to establish a LIG4-deficient HAP1 population (HAP LIG4 KOp), which circumvents clonal expansion (Cortazar & Jagannathan, 2025). In this line, sequenced reads from gDNA predominantly originate from HDR events, 74.24% ± 2.61 compared to 7.50% ± 1.13 in parent cells (Fig. 1C; Fig. S1A). No contamination from the plasmid library was detected. Co-transfection of the donor plasmid library with a non-targeting sgRNA plasmid did not yield detectable HDR events from gDNA (Fig. 1C; Fig. S1A).

Using the HAP1 LIG4 KOp cells, we performed triplicate SGE experiments targeting LMNA exons and quantified variant expression levels by normalizing RNA read counts to gDNA frequencies. After stringent filtering (see Methods), we obtained robust measurements of mRNA expression for 722 PTC and 211 SNV *LMNA* variants under both DMSO and SMG1i conditions (86.6% of all intended mutations), with high replicate correlation (Fig. S1B). As expected, PTC variants in general caused depletion of the LMNA mRNA, while SNVs did not (Fig. 1D, E). The correlation between DMSO and SMG1i conditions was lower for PTCs (r = 0.62-0.75) compared to SNVs (r = 0.95), consistent with NMD selectively targeting PTC-containing transcripts.

Indeed, SMG1i rescued PTC expression but did not affect SNV levels (Fig. S4B). SGE-treated cell populations did not exhibit any substantial phenotypic change, consistent with *LMNA* being non-essential (Harborth et al., 2001; Rober et al., 1989). *LMNA* null mice develop normally and only die 1-2 months after birth due to muscular dystrophy (Sullivan et al., 1999). Together, these results demonstrate that SGE of LMNA provides a robust system for systematic and quantitative analysis of NMD activity.

### A PTC position effect suggests a short-range influence of the last EJC on NMD

We first evaluated whether our SGE strategy can capture the expected transition in NMD activity (active to inactive) dictated by the 50-nt rule. To this end, we targeted the 3′ end of exon 11, which encompasses the 50-nt rule boundary. In addition to assessing NMD efficiency, we examined the effects of PTCs on transcription and splicing to obtain a more comprehensive view of their impact on LMNA mRNA.

Following SGE targeting of exon 11, a fraction of the cell population was used to assess NMD efficiency, while the remaining fraction was subjected to pulse labelling of nascent transcripts with 5-bromouridine, followed by immunoprecipitation (Bru-IP) and sequencing (Fig. 2A) (Paulsen et al., 2014). We first validated the protocol for analysis of nascent transcripts as follows. BrU-IP followed by qRT-PCR confirmed enrichment of pre-mRNA signal including intergenic regions from the *GAPDH* gene. There was minimal mature RNA contamination as indicated by comparable intronic and exonic signals (Fig S2A, B) and specific enrichment of a BrU-labeled spike-in control. The detection of a relatively small fraction of exon–exon junctions is consistent with co-transcriptional splicing in human genes (Shenasa & Bentley, 2023). A multiplex PCR-gel electrophoresis assay further verified enrichment of nascent LMNA transcripts (Fig. 2B). *LMNA* encodes two mRNA isoforms, Lamin A and Lamin C. Lamin A differs from Lamin C by splicing at exon 10 and contains two unique 3’ end exons, 11 and 12. We therefore performed targeted sequencing by priming cDNA synthesis in exon 12 to analyze spliced Lamin A mRNA, and in intron 11 to capture unspliced pre-mRNA.

**Figure 2.**
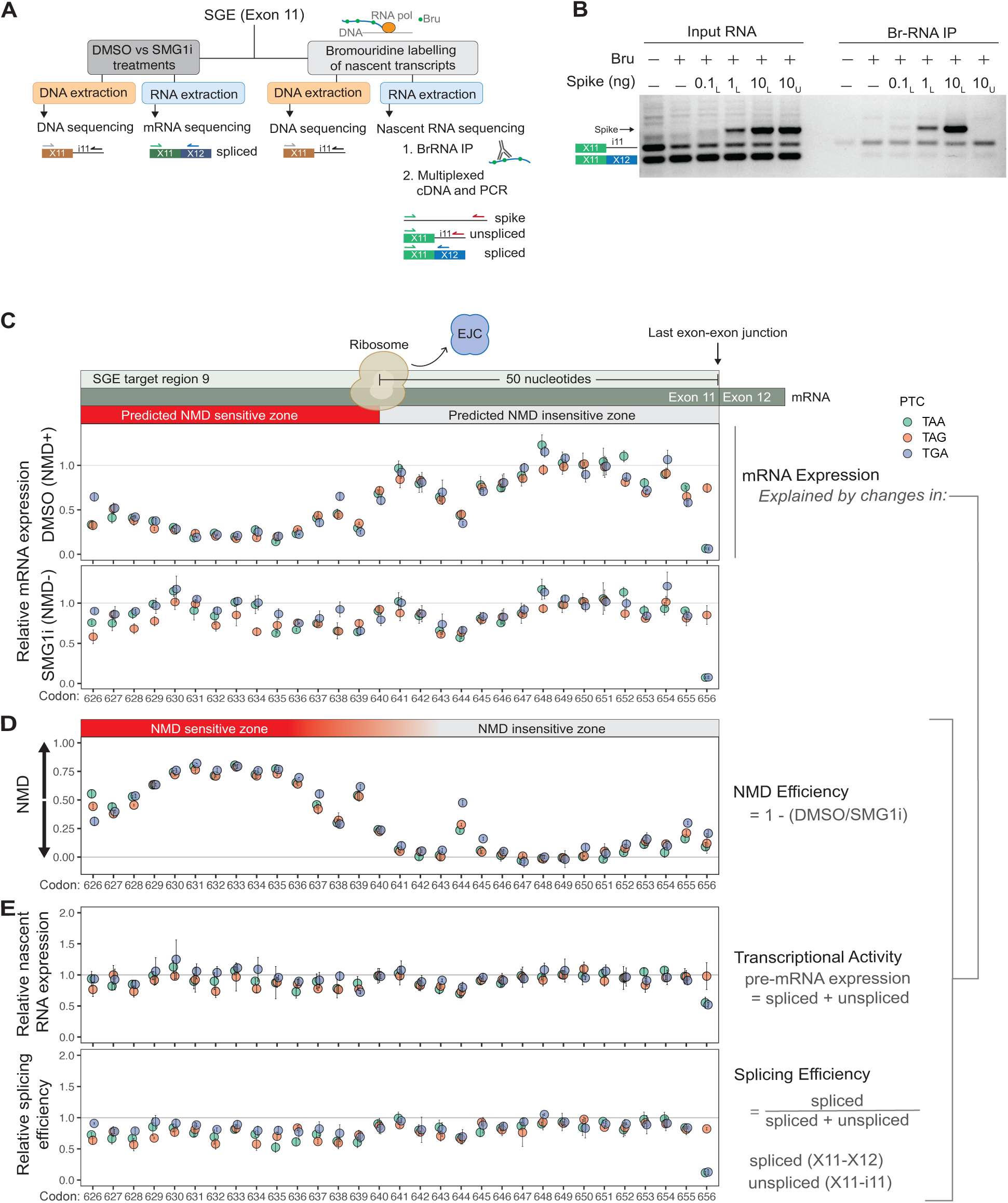
PTC position effect suggests a short-range influence of the last EJC on NMD. **(A)** SGE-treated cells were treated with SMG1i or DMSO and incubated with bromouridine (BrU) for metabolic labelling of nascent transcripts. Targeted sequencing libraries were prepared using multiplex PCR to detect exon 11–exon 12 junctions (X11–X12) and the unspliced exon 11–intron 11 boundary (X11–i11) from isolated nascent transcripts. **(B)** BrU-labelled RNA (Br-RNA) immunoprecipitation (IP), followed by RT-PCR and gel electrophoresis, showing enrichment of unspliced and spliced LMNA nascent transcripts. An *in vitro*–transcribed RNA spike-in was added in increasing amounts to the input sample and was either BrU-labelled (L) or unlabelled (U). Note the specific enrichment in the IP only when the spike-in is BrU-labelled. **(C)** mRNA expression of *LMNA* variants containing a single PTC at the indicated codon position, relative to the synonymous sequence control, under DMSO or SMG1i conditions. **(D)** NMD efficiency of PTC variants. **(E)** Transcriptional activity and splicing efficiency relative to the synonymous sequence control.

For analysis of mRNA expression levels and NMD efficiency, variant frequencies were quantified by targeted sequencing of gDNA and total RNA. As expected from the 50-nt rule, PTCs upstream of codon 640 caused stronger reduction in mRNA levels than those downstream, and their expression was rescued by SMG1i (Fig. 2C). SNVs which do not incorporate PTCs neither reduced mRNA levels nor responded to NMD inhibition (Fig. S2C). Cycloheximide (CHX), a translation inhibitor, similarly rescues the expression of PTC variants in the predicted NMD sensitive zone, confirming the dependence of NMD on translation (Fig. S2D). CHX treatment was not as efficient at rescuing PTC-containing variants as SMG1i. This difference may be explained by the fact that cycloheximide interferes with NMD indirectly by inhibiting translation elongation, likely introducing secondary effects that also impact mRNA stability, whereas SMG1 inhibition directly blocks UPF1 phosphorylation, a central step in the NMD pathway. We calculated NMD efficiency as the fraction of mRNA degraded by NMD. The expression level after NMD inhibition with SMG1i (Fig. 2D) or CHX (Fig. S2D) represents the total population available to NMD, while expression in DMSO condition represents the fraction evading NMD. SMG1i and CHX yielded nearly identical NMD efficiency profiles (Fig. 2D, Fig. S3A).

Strikingly, plotting NMD efficiency around the 50 nt boundary revealed a smooth, position-dependent effect rather than a sharp transition from strong to no NMD (Fig. 2D). NMD activity peaked at codon 633 (approximately 71 nt upstream of the last exon-exon junction) and declined gradually as the PTC moved away from this point in the 5’ or 3’ direction. This profile suggests that an optimal distance between the terminating ribosome and the last EJC downstream is the major determinant of NMD efficiency in the penultimate exon. This distance-dependence may explain why long exons (>400 nt) are associated with reduced NMD activity (the “long exon” rule; (Lindeboom et al., 2016)), where a greater proportion of the exon lies far from the nearest EJC (Hoek et al., 2019). PTC identity is not a major source of NMD variability in this region, with only a few codon positions (626, 644 and 645) showing divergence among TAA, TAG, and TGA. Notably, intermediate levels of NMD activity can also be detected within the predicted NMD insensitive zone at codon 644, consistent with previous reports indicating that an EJC is not an absolute requirement for NMD (Buhler et al., 2004; Fang et al., 2013).

To determine whether the observed NMD profile is specific to HAP1 cells, or intrinsic to the LMNA mRNA, we performed the same experiment in wild-type HEK293 cells (Fig. S2E; Fig. S3B). Without *LIG4* inactivation, HDR frequency was lower (2.65% ± 0.10 reads) resulting in higher noise in the NMD efficiency measurements. Nevertheless, all intended variants were integrated successfully, and the NMD activity profile closely reflects the behavior seen in HAP1 cells (Fig. S3A). Thus, the position-dependent NMD behavior around the 50-nt rule boundary is a conserved property of the LMNA transcript.

Beyond NMD, PTCs also impact transcriptional activity and splicing efficiency in a position-specific manner (Fig. 2E). Together, these effects explain the overall steady state mRNA levels. For example, PTCs at codon 656, the last codon of exon 11, dramatically reduced mRNA expression through reduced transcriptional activity and splicing efficiency, but only when the PTC introduced removed the canonical “G” nucleotide in the −1 position of the 5’ splice site consensus (i.e., TAA and TGA show reduced mRNA, but not TA<u>G)</u> (Parker et al., 2025).

Therefore, NMD inhibition does not fully rescue the expression of TAA and TAG variants (Fig. 2C) as it cannot rescue the splicing defect. Although PTCs at codon 656 caused a pronounced splicing defect, many others (37 out of 93) also reduced splicing efficiency by approximately 20–40% (Fig. 2E; Fig. S3C). These results highlight a critical limitation of inferring NMD activity from steady-state mRNA levels alone: given the potential contributions of transcription and splicing defects to mRNA expression, direct measurement of the mRNA fraction targeted by NMD is essential for accurate calculation of NMD activity. Importantly, our results demonstrate that NMD activity is highly variable immediately upstream of the 50-nt boundary and that its efficiency can be largely accounted for by the distance between the prematurely terminating ribosome and the last exon–junction complex.

### PTC position dictates NMD strength and uncovers activation–deactivation gradients along the mRNA

We next asked how PTC position within the LMNA transcript influences mRNA expression and NMD activity. Building on our exon 11 analysis, we extended SGE to exons 1, 4, 7, and 10, and normalized the expression of each PTC variant to the median expression of the SNV controls within each experiment. SNV-based normalization did not alter the overall expression or NMD activity profile but improved consistency and alignment across adjacent SGE target regions (Fig. 3A, C; Fig. S4A-C).

**Figure 3.**
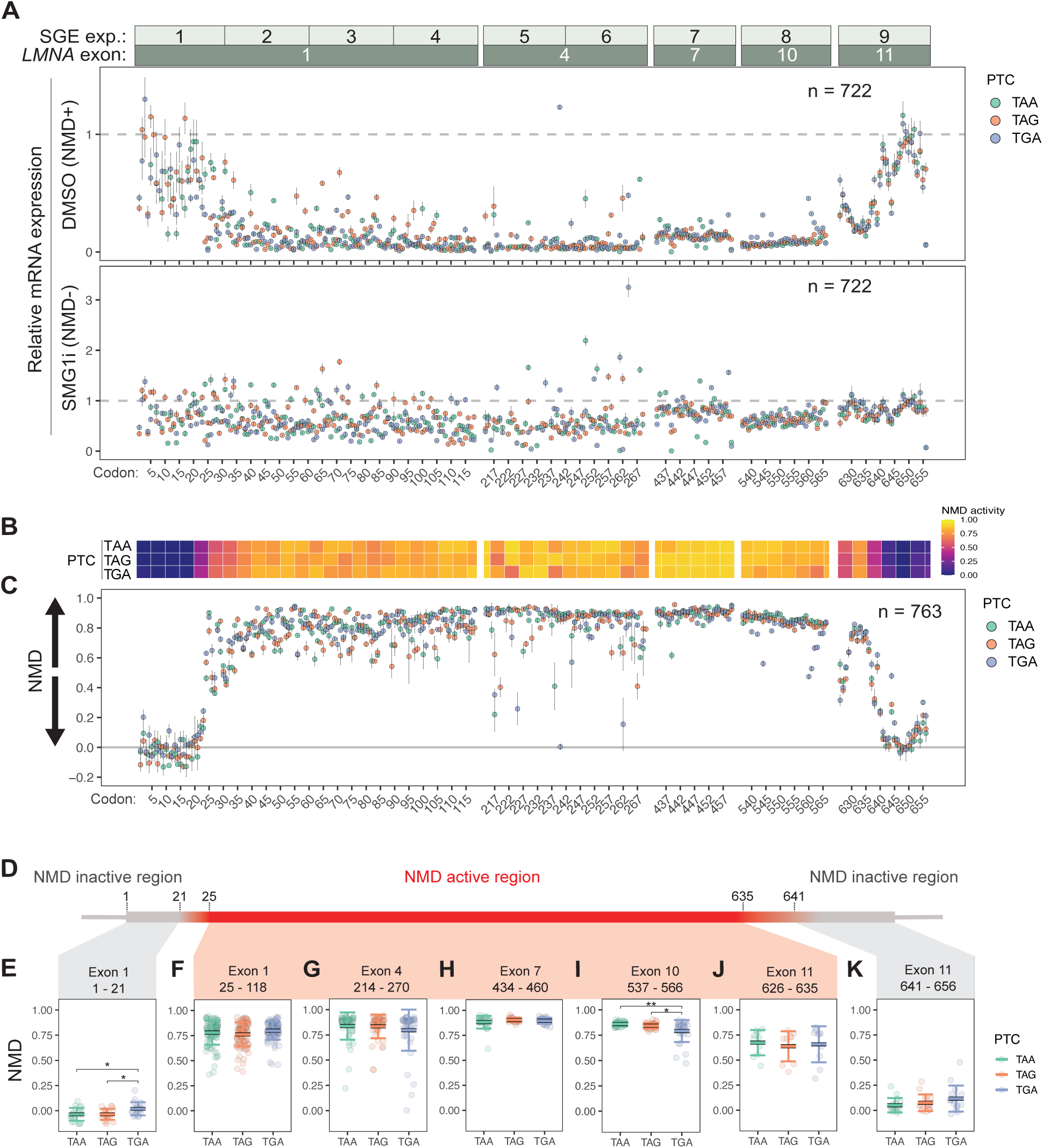
PTC position dictates NMD strength and uncovers activation–deactivation gradients along the mRNA. **(A)** mRNA expression levels of PTC variants relative to the synonymous sequence context control under basal conditions (DMSO) or following inhibition of NMD with SMG1i. **(B)** Heat map of mean NMD efficiency calculated in 5-codon windows for each stop codon type (TAA, TAG, TGA). **(C)** NMD efficiency of PTC *LMNA* variants across the mRNA. **(D)** Schematic map of NMD-inactive and NMD-active regions along the LMNA mRNA. NMD transition zones occur between codons 21–25 and 635–641. **(E–K)** Mean NMD efficiency across exons. A Kruskal–Wallis test indicated significant differences among stop codon types. Post hoc pairwise Wilcoxon rank-sum tests with Benjamini–Hochberg correction showed that TGA differed significantly from both TAA (adjusted p = 0.025) and TAG (adjusted p = 0.014) in exon 1 (panel **E)**, and that differences were observed between TGA and both TAG (adjusted p = 0.042) and TAA (adjusted p = 0.004) in exon 10 (panel **I)**. Note that calculation of mRNA expression levels requires variants to pass quality filters in both cDNA and gDNA libraries, whereas NMD efficiency calculations require variants to pass filters only in the cDNA libraries (see Methods). Consequently, more variants were excluded from the mRNA expression analysis (n = 722) than from the NMD efficiency analysis (n = 763).

Systematic integration of PTCs across the transcript revealed a striking U-shaped RNA expression profile that mirrors the inverted U-shaped profile of NMD activity: mRNA levels were low throughout internal exons but preserved at both ends of the ORF (Fig. 3A). SMG1i treatment rescued expression of most internal PTC variants, confirming that NMD is the primary driver of this pattern. However, a significant subset of PTCs remained very lowly expressed even after NMD inhibition, demonstrating NMD-independent effects on mRNA abundance, potentially involving transcriptional or splicing defects, as observed for exon 11 (Fig. 2E).

Conversely, some PTC variants in exon 1, 4, and 7 exceeded wild-type expression levels upon SMG1i treatment (Fig. 3A; points above the dashed SMG1i line), possibly reflecting nonsense-induced transcriptional adaptation (reviewed in (Kontarakis & Stainier, 2020)).

To visualize NMD along the transcript, we first calculated mean activity in 5-codon windows for each stop codon type (TAA, TAG, and TGA) (Fig. 3B). The resulting heatmap reveals a clear positional pattern, where NMD activity is uniformly high (red/orange) across internal exons but drops sharply (blue) at both 5’ and 3’ ends of the ORF. The three stop codons follow the same general trend. Single-codon resolution analysis confirmed this pattern, showing an inverted U-shaped NMD profile (complementary to the U-shaped expression profile in Fig. 3A), where the magnitude of NMD can be explained by the location of the PTC along the transcript (Fig. 3C). The transitions between active and inactive zones are gradual rather than abrupt, highlighting the quantitative relationship between PTC position and the magnitude of NMD activation.

We defined the NMD-active region as the interval between the activation and deactivation boundaries, codons 25 to 635, spanning ∼75 nt downstream of the start codon to ∼66 nt upstream of the normal termination codon (Fig. 3D). Within this region, mean NMD efficiency was 0.806 ± 0.125, demonstrating that NMD strongly degrades the vast majority of PTC-containing transcripts. Notably, NMD activity showed no global 5’ or 3’ bias across internal exons, suggesting that the total number of EJCs upstream or downstream of the PTC does not substantially influence NMD on this transcript.

While PTC position is the dominant determinant of NMD, local variation is also evident at discrete codon positions. Codon-specific drops in NMD are strongly influenced by PTC identity, as reflected by variation among TAA, TAG, and TGA within same codon positions (Fig. 3C). Notably, there are no global differences in mean NMD activity across the three PTC types within the NMD active region, but dispersion across PTC variants is most pronounced near the 5′ end of the transcript in exons 1 and 4 (SD = 0.11, 0.16), compared to exons 7 and 10 (SD = 0.04, 0.07) (Fig. 3E-K). These findings indicate that both the PTC position and the stop codon identity modulate NMD, and that their effects are dependent on the specific exon.

### NMD is inactive against start-proximal PTCs and activates in a sharp, position-dependent gradient

Previous studies have shown that NMD is inefficient in targeting start-proximal PTCs, leading to the formulation of the “200-nt rule” (Lindeboom et al., 2016). This rule posits that PTCs within 200 nt of the start codon are less efficient in triggering NMD. Our data in *LMNA* reveal a sharper picture. PTCs within the first 21 codons fail to trigger NMD, with only a few variants showing weak activity (Fig. 4A). NMD then activates gradually over a very short span of four codons (positions 22–25). Beyond this activation zone, NMD fluctuates across adjacent codons but is generally robust (Fig. 3A, B).

**Figure 4.**
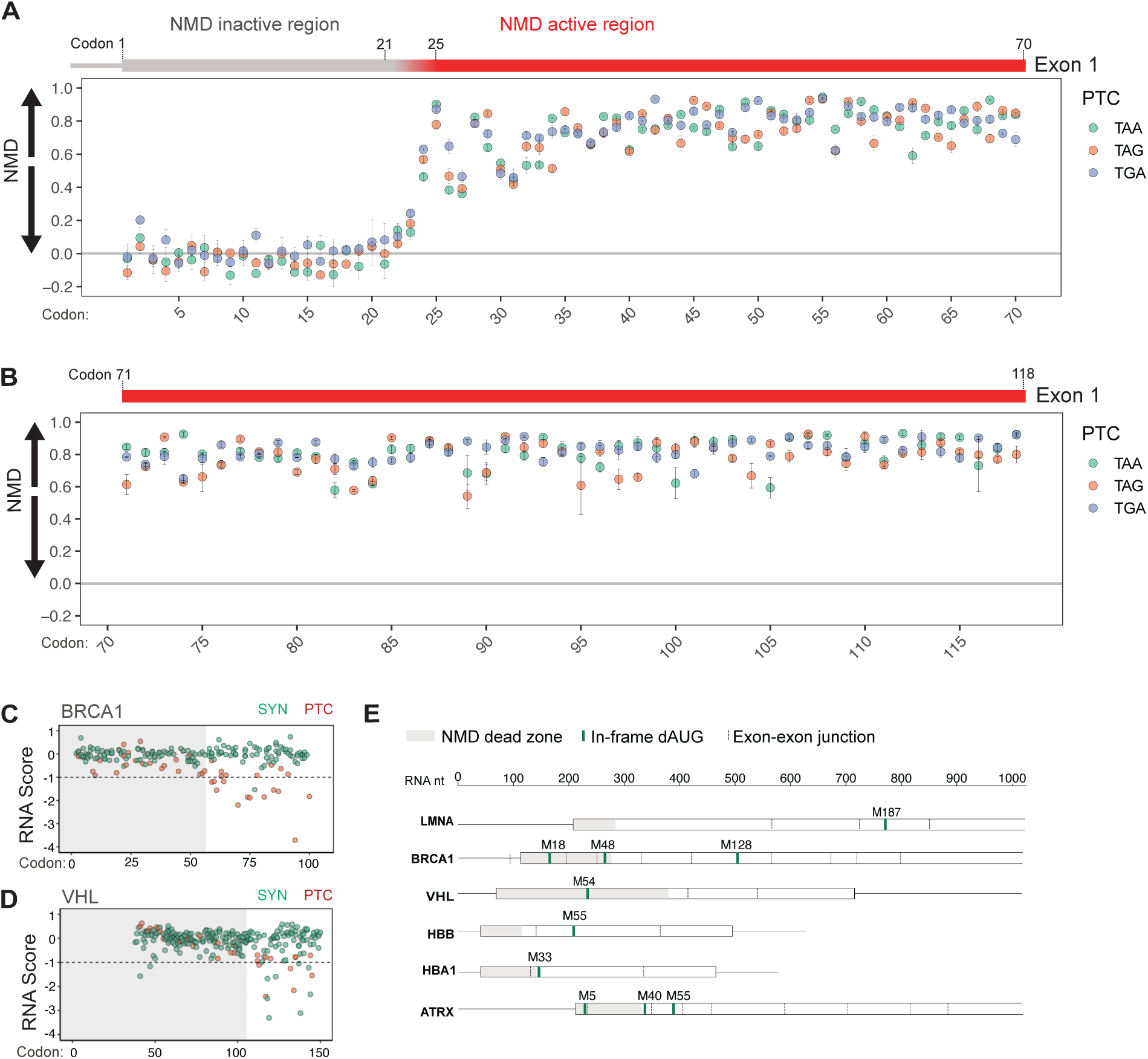
NMD is inactive against start-proximal PTCs and activates in a sharp, position-dependent gradient. NMD efficiency along the full length of LMNA exon 1 for PTC variants located in the 5′ half **(A)** and 3′ half **(B)**. **(C-D)** RNA expression scores for synonymous (SYN, green) and PTC (red) variants across the start-proximal region of *BRCA1* (Findlay et al., 2018) and *VHL* (Buckley et al., 2024). Gray shading indicates the NMD dead zone. **(E)** Schematic comparison of start-proximal NMD-inactive regions across seven genes. Boundaries for LMNA are from this study; boundaries for BRCA1 and VHL are derived from published saturation genome editing data; (Buckley et al., 2024; Findlay et al., 2018) boundaries for *HBB*, *HBA1*, and *ATRX* are based on published reports (Danckwardt et al., 2002; Howard et al., 2004; Inacio et al., 2004; Neu-Yilik et al., 2011; Peixeiro et al., 2012; Pereira et al., 2015; Silva et al., 2006). Downstream in-frame start codons were annotated using NetStart 2.0 (Nielsen et al., 2025).

To determine whether this start-proximal NMD inactive zone is specific to LMNA or a general feature, we analyzed published saturation genome editing datasets from BRCA1 (Findlay et al., 2018) and VHL (Buckley et al., 2024). Note that in these experiments, PTCs were only sparsely sampled, and NMD activity was not directly measured, but inferred from relative RNA abundance of PTCs compared to synonymous variants. In BRCA1, PTCs throughout the first ∼50 codons show minimal mRNA depletion compared to synonymous controls, followed by sharp depletion of PTC variants (Fig. 4C). Similarly, VHL displays a discrete NMD-inactive zone at its 5′ end with an abrupt transition to mRNA depletion (Fig. 4D). These data demonstrate that start-proximal NMD boundaries are a conserved feature across genes, though the precise position of the boundary varies.

To explore the basis of the boundary position further, we compared NMD-inactive regions across seven genes for which start-proximal NMD data are available (Fig. 4E). The length of the NMD dead zone varies substantially, ranging from ∼21-23 codons in LMNA and HBB to ∼100 codons in VHL. Notably, the presence and position of in-frame downstream AUGs does not predict the boundary. *LMNA* lacks in-frame AUGs within its first coding exon, yet shows a boundary similar to HBB, which contains a downstream AUG in close proximity. Conversely, BRCA1 has in-frame AUGs within its NMD-inactive region (M18, M48) but still shows NMD resistance extending to codon ∼50. These observations suggest that while reinitiation may contribute to NMD evasion in some contexts, it does not explain the start-proximal boundary.

Two mechanisms have been proposed to explain NMD suppression at start-proximal PTCs: translation reinitiation downstream of the PTC and PABPC1-stimulated efficient translation termination in the mRNA circularization context (Inacio et al., 2004; Lindeboom et al., 2016; Neu-Yilik et al., 2011). In the LMNA transcript, no in-frame AUG codons exist downstream of the start site within exon 1 or exon 2, excluding canonical reinitiation. Alternative CUG codons are present (at positions 21, 29, 35, 52, 59 and 102) but in poor Kozak contexts, making non-canonical reinitiation an unlikely explanation. Our data instead support the termination efficiency model: within the NMD inactive region, TGA variants show significantly higher residual NMD activity than TAA or TAG variants (Wilcoxon rank-sum test with Benjamini–Hochberg correction: TGA vs. TAA, p = 0.025; TGA vs. TAG, p = 0.014) (Fig. 3E). Because TGA is the least efficient termination codon (Manuvakhova et al., 2000; Wangen & Green, 2020), reduced PABPC1-mediated termination efficiency at TGA may increase the opportunity for activation of NMD. Taken together, these observations support a model in which efficient termination, potentially facilitated by PABPC1 proximity, suppresses NMD at start-proximal PTCs.

The mechanism underlying start-proximal NMD inactivation and what determines their gene-specific boundaries remains unclear. Boundary position is likely determined by the physical distance between the terminating ribosome and PABPC1 in the closed-loop mRNA conformation and further influenced by reinitiation potential or other transcript-specific features. Regardless of mechanism, our multi-gene analysis establishes that start-proximal NMD inactive boundaries are a general feature of mammalian transcripts.

### Local sequence contexts modulate NMD efficiency in an exon-dependent manner

We next asked whether local sequence context surrounding the PTC influences NMD activity if we only consider the NMD-active regions of the transcript. We systematically examined nucleotide identity at positions −1, −2, −3 and +4, +5, +6 flanking the PTC across all exons (Fig. 5A-F). Of these positions, only −1 and +4 show consistent effects on NMD. An A at the −1 position is associated with reduced variability and robust NMD activation across all exons containing this context (exon 1, 4 and 10) (Fig. S5A-E), suggesting this context limits the influence of NMD-attenuating mechanisms. However, the most striking effects occur at the +4 position, where NMD activity varies substantially depending on both downstream nucleotide and the PTC identity.

**Figure 5.**
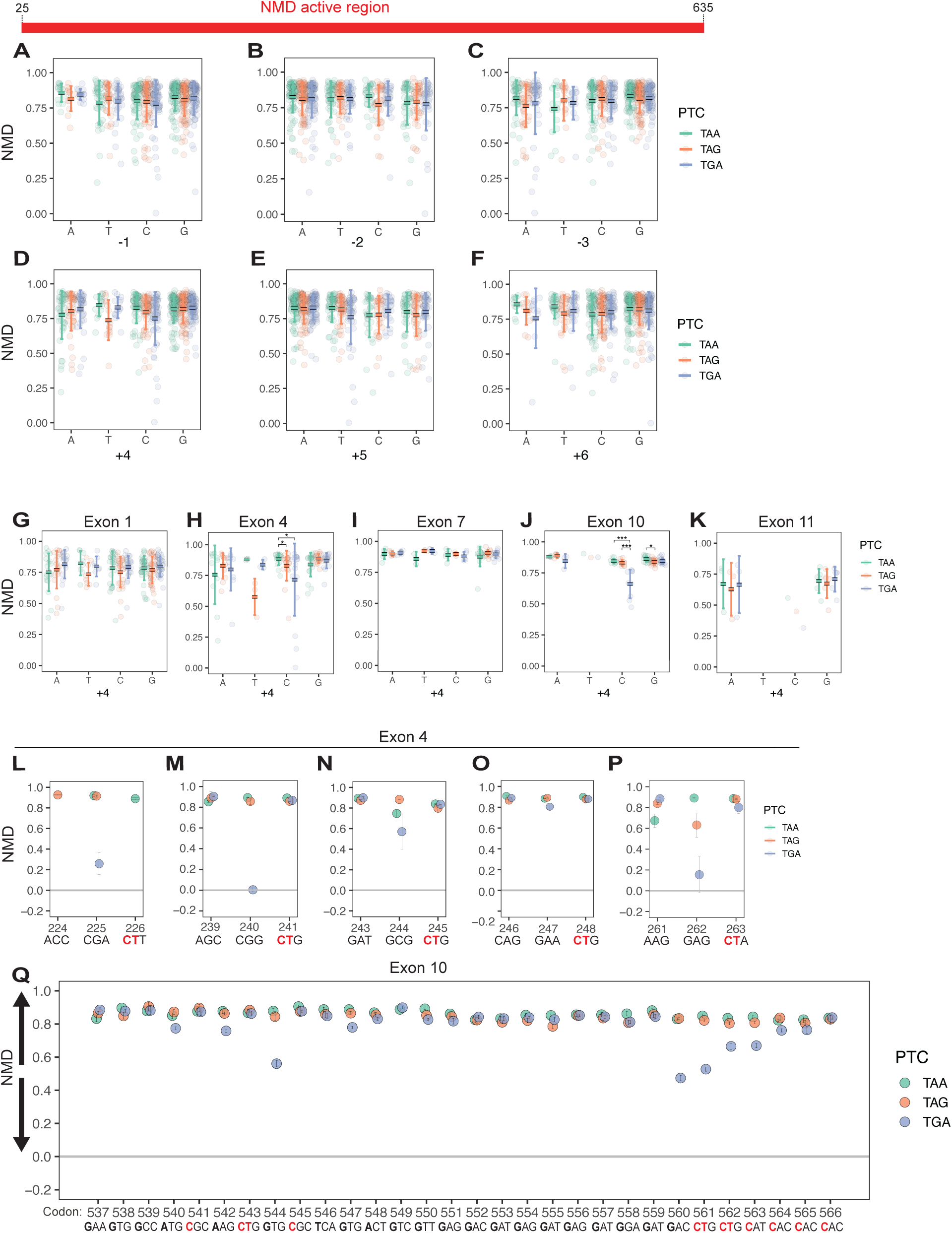
Local sequence contexts modulate NMD efficiency in an exon-dependent manner. (A–F) Mean NMD efficiency as a function of the nucleotide identity at positions −3, −2, −1, +4, +5, and +6 relative to the PTC within the NMD-active region. **(G–K)** Same as in (A–F), analyzed separately for each exon in the NMD-active region. **(L–P)** NMD variation at specific codon positions marked by local drops in NMD efficiency within the TGA-CT sequence context in exon 4. **(Q)** Same as in (L–P), shown across the full length of exon 10. The control sequence context is shown beneath panels L–Q. Post hoc pairwise Wilcoxon rank-sum tests with Benjamini–Hochberg correction showed that TAA differed significantly from both TAG and TGA (adjusted p = 0.039 for both comparisons) in exon 4 (panel **H**), whereas the same analysis showed that TGA differed significantly from both TAA and TAG (adjusted p = 6.2 × 10⁻⁵ for both comparisons in exon 10 (panel J).

C at the +4 position of TGA, a known readthrough promoting context, strongly correlates with reduced NMD efficiency, particularly in exons 4, 10, and 11 (Fig. 5D, G-K). At positions with a +4 C, the relative NMD strength among the stop codons becomes TAA>TAG>TGA (Fig. 5D, H, J, K), inversely matching the established hierarchy of readthrough permissiveness among the stop codons (TGA>TAG>TAA). This supports a model where translation readthrough may suppress NMD by allowing the ribosome to move past the PTC to eject downstream EJCs. A similar but weaker pattern emerges for TAG followed by +4 T in exons 1 and 4 (Fig. 5D, G, H), although the small number of variants in these contexts limits the statistical power to establish additional significant associations. Interestingly, a T at the +5 position, also emerges as a context associated with outliers of weaker NMD activity (Fig. 5E, Fig. S5F-J), specifically in exons 4 and 10 (Fig. S5G, I). Notably, the dispersion among PTCs is greater in exons 1 and 4 compared to downstream exons, suggesting a 5’ bias in the influence of the PTC sequence context on NMD.

To examine these effects at single-codon resolution, we focused on exons 4 and 10 (Fig 5. G-K). In exon 4, five out of six TGA-CT contexts show substantially reduced NMD activity (Fig.5L-P; Fig. S5G), showing an influence of the +5 T nucleotide (Fig S5G). Strikingly, the TGA-CT context at codon 240 is entirely NMD-resistant, representing the weakest NMD context in the entire NMD-active region. In exon 10, TGA results in reduced NMD efficiency only when followed by a +4 C, and TGA-CT is among the contexts that most strongly inhibit NMD at codon 560 and 561 (Fig. 5Q; Fig. S5I). While translation readthrough remains to be demonstrated on the LMNA mRNA, TGA-CT is a conserved and efficient readthrough motif across diverse species (Stiebler et al., 2014), strongly implicating translation readthrough as the mechanism underlying local NMD evasion at these positions.

Together, these findings reveal that NMD variation within the active regions occurs at discrete codon positions rather than gradually across neighboring sites. Because this variation depends on stop codon identity and the immediate downstream nucleotide (features recognized within the ribosomal A site during translation termination), we propose that differences in stop codon decoding by eRF1 underlie these codon-specific effects on NMD.

### NMD variability on the LMNA mRNA mirrors NMD behavior across the human transcriptome

To determine whether the positional patterns of NMD activity observed in LMNA reflect general principles across the transcriptome, we analyzed the allele specific expression of 7,141 naturally occurring heterozygous stop gain variants (Fig. 6A). Using paired whole genome sequencing and RNA-seq datasets from the NHLBI Trans-Omics for Precision Medicine (TOPMed) program, we calculated a normalized NMD score as the ratio of reference allele read counts to total read counts, where higher values indicate stronger NMD-mediated degradation of the PTC-containing allele.

**Figure 6.**
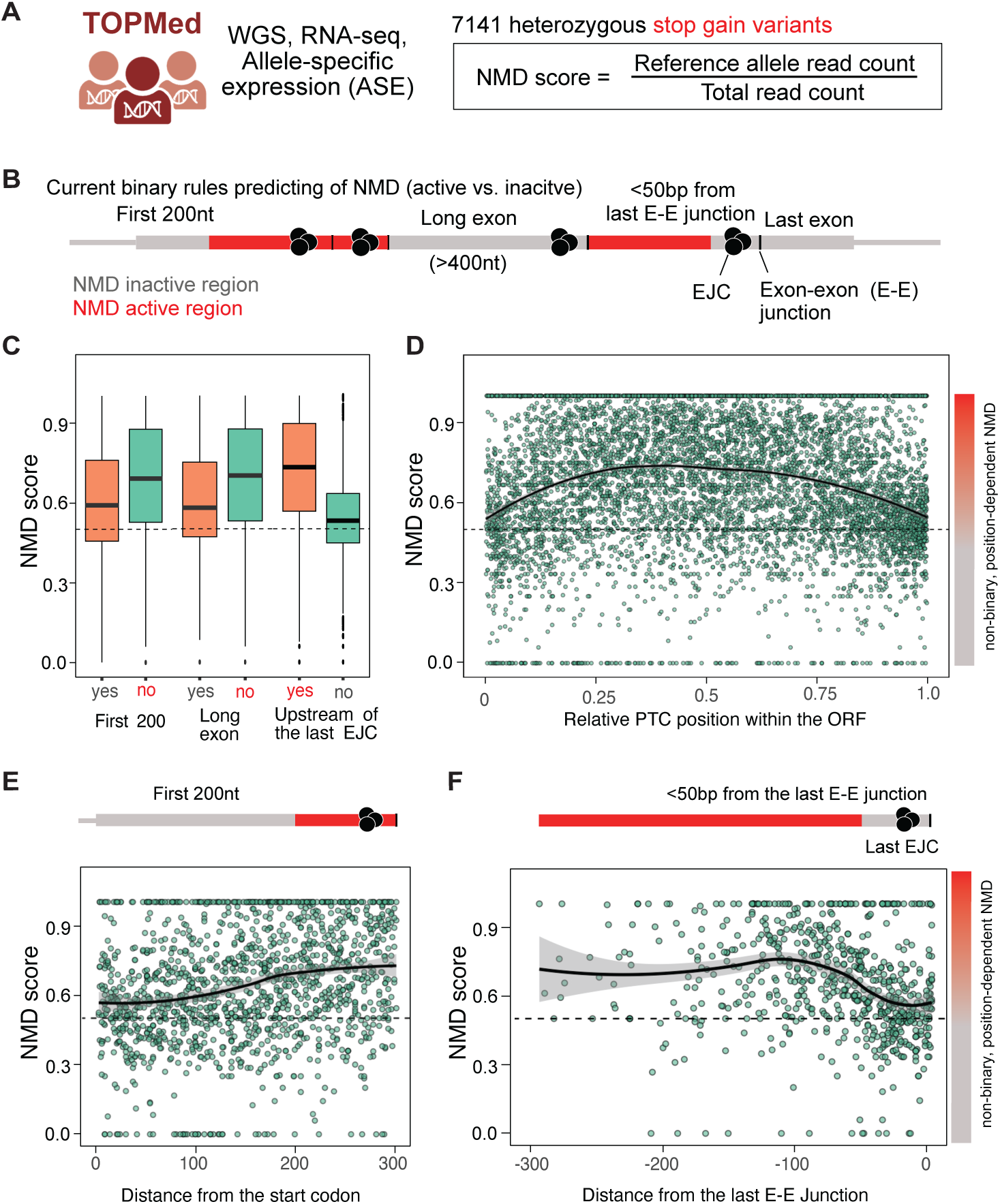
NMD variability on the LMNA mRNA mirrors NMD behavior across the human transcriptome. **(A)** Calculation of the NMD score for TOPMed nonsense variants. **(B)** Graphical illustration of current NMD rules. **(C)** NMD score categorized for nonsense variants located within the first 200 nt, in long exons, or upstream of the last exon junction complex (EJC). **(D)** NMD score for variants at relative positions along the gene ORF. **(E)** Same as in (D) but plotted as a function of the distance of each nonsense variant relative to the translation start site. **(F)** Same as in (D) but plotted as a function of the distance of each nonsense variant relative to the downstream last exon–exon junction.

Current models describe NMD behavior using binary positional rules that are deterministic: start-proximal PTCs in the first ∼200 nt of the ORF, PTCs in long exons (>400 nt), and PTCs in the last exon or within 50 nt of the last exon-exon junction evade NMD (Fig. 6B; “NMD inactive”), while PTCs in the rest of the transcript undergo NMD (Fig. 6B; “NMD active”). We first confirmed that each of these rules holds at the population level in our dataset (Fig. 6C): PTCs in the first 200 nt show a lower NMD score than those outside this region; PTCs in long exons exhibit weaker NMD compared to shorter exons; and PTCs upstream of the 50-nt boundary trigger stronger decay than those downstream of this boundary. Although these categorical trends are qualitatively consistent with the binary rules, plotting NMD score as a function of relative PTC position within the ORF reveals substantial quantitative variability that binary rules cannot capture (Fig. 6D). The data show a broad inverted U-shaped profile, with reduced NMD at both the 5′ and 3′ ends of the ORF and higher score in the central region, qualitatively matching the pattern observed in LMNA (Fig. 3C). However, the variability around this trend is considerable. Many PTCs predicted to trigger efficient NMD by binary rules show weak decay, while others in “NMD-inactive” zones are efficiently degraded. This variability underscores the limitations of applying binary filters to predict NMD outcomes for individual variants.

At the 5′ end of the ORF, the population-level data show a gradual increase in NMD score over the first ∼200 nt (Fig. 6E), in apparent contrast to the sharp 4-codon activation boundary observed in LMNA (Fig. 4A). However, our multi-gene analysis reveals that start-proximal NMD-inactive zones are a general feature but have different boundaries in different genes (Fig. 4E). The gradual population-level profile therefore likely reflects the superposition of gene-specific boundaries at varying positions, which average to a smooth transition when aggregated across thousands of transcripts (Lindeboom et al., 2016).

In contrast, the 3′ boundary of NMD activity shows remarkable consistency between single-gene and transcriptome-level analyses. As PTCs approach the last exon-exon junction, NMD score declines in a smooth, distance-dependent manner (Fig. 6F), precisely recapitulating the quantitative PTC-EJC relationship we characterized in LMNA exon 11 (Fig. 2D). This concordance supports a model in which NMD efficiency at the 3′ end of the ORF is governed by the spatial relationship between the terminating ribosome and the downstream EJC, rather than a binary 50-nt threshold.

Together, these transcriptome-wide analyses validate and extend our LMNA findings. While binary positional rules capture broad trends, they fail to account for the substantial variability in NMD outcomes across individual PTCs. The gradual NMD activation of start-proximal PTCs at the population level likely reflects the averaging of sharp, gene-specific boundaries. The 3′ boundary, in contrast, shows a conserved quantitative relationship with PTC-EJC distance that is consistent between single-gene and population-level data. These observations highlight the value of single-gene, codon-resolution mapping for uncovering quantitative predictors of NMD activity and their mechanistic underpinnings.

## Discussion

Using a saturation genome editing–based approach, we systematically introduced 722 PTCs and 211 SNV control variants across the LMNA coding sequence and directly quantified their effects on mRNA expression and NMD. This quantitative, high-resolution map reveals that NMD is not governed by simple binary rules but instead varies substantially with position and sequence context, highlighting a far more nuanced regulatory landscape than previously appreciated.

A striking feature of our codon-resolution NMD map is the inverted U-shaped profile of NMD activity across the length of the LMNA transcript. This profile defines three discrete zones: a start-proximal “dead zone” where PTCs fail to trigger NMD (codons 1-21); sharp activation of NMD over codons 22-25 and maintenance of high efficiency (mean of 0.81 ± 0.13) over most of the internal exons, and a quantitative gradient of NMD activity decline in the penultimate exon rather than a sharp binary transition predicted by the classical 50-nt rule. This Gaussian profile of NMD upstream of the 50-nt boundary (Fig. 2D) is also evident at the transcriptome level (Fig. 6F), highlighting its broad applicability across transcripts. Analysis of published saturation genome editing data from BRCA1 and VHL, together with literature reports from additional genes, confirms that sharp start-proximal NMD boundaries are a general feature of mammalian transcripts, though the position of this boundary varies across genes (Fig. 4E). These distinct regulatory zones of NMD activity along the transcript provide a quantitative framework for how PTC position governs NMD outcomes and suggests that distinct molecular mechanisms influence NMD at each of these zones.

At the 5’ end of the LMNA mRNA, we captured a sharp NMD activation that spans only four codons (22-25), following a start-proximal region that is refractory to NMD. This behavior contrasts previous RNA-seq meta-analysis results showing a gradual ∼200 nt transition near the start codon (Lindeboom et al., 2016), as well as our own transcriptome-wide analyses (Fig. 6E). However, our analysis of published BRCA1 and VHL saturation genome editing data reveals similarly sharp NMD activation boundaries, though at more distal positions (∼codon 50 in BRCA1, ∼codon 100 in VHL) than in LMNA or β-globin (∼codon 22–25) (Fig. 4C–E). These observations suggest that sharp start-proximal NMD boundaries are a general feature of mammalian transcripts, with gene-specific positions that are obscured when aggregated at the population level.

What determines the position of the NMD activation boundary in each gene remains unclear. Translation re-initiation is the mechanism most frequently invoked to explain NMD resistance of start-proximal PTCs. However, comparison across several genes reveals that the presence and proximity of downstream AUGs is neither necessary nor sufficient to establish this boundary (Fig. 4E). BRCA1 contains in-frame AUGs within its NMD-inactive region yet still shows NMD resistance extending to codon ∼50, while LMNA lacks such AUGs but has a boundary at codon 22–25. Instead, the residual NMD activity observed specifically for TGA-containing variants, the stop codon with the weakest termination efficiency, suggests that NMD inactivation at start-proximal PTCs may be the result of efficient translation termination stimulated by PABPC1 (Amrani et al., 2008; Peixeiro et al., 2012; Silva et al., 2008). This model is supported by evidence that PABPC1 competes with UPF1 for direct binding to the eukaryotic release factor eRF3 (Singh et al., 2008), and by the well-established notion that a closed-loop mRNA conformation promotes efficient translation initiation by bringing the poly(A) tail into close proximity with the 5′ end (Amrani et al., 2008; Gallie, 1991). The sharp activation of NMD around codon 22 on both the LMNA (Fig. 4A) and β-globin transcripts (Inacio et al., 2004; Neu-Yilik et al., 2011) may therefore mark a point in early elongation at which spatial communication between the terminating ribosome and PABPC1 is lost. Given the different lengths of the start-proximal NMD-inactive zone across several transcripts (Fig. 4E), the point of transition may occur within 22 to 100 codons from the translation start site, though additional genes need to be tested to establish generality. In other transcripts, this boundary may be extended by reinitiation, alternative mRNA conformations, or other transcript-specific features.

Interestingly, the length of the start-proximal NMD inactivity zone coincides with the short median length (∼22 codons) of uORFs, which occur in over 50% of human transcripts and yet only a subset trigger NMD (May et al., 2023; Stockklausner et al., 2006). Whether this reflects selective pressure on uORF length to avoid triggering NMD, or whether NMD suppression at start-proximal positions evolved to tolerate uORF translation, remains to be determined.

On the other end of the LMNA transcript, the Gaussian profile of NMD activity obtained around the 50-nt boundary suggests that the influence of the last exon junction complex (EJC) is highly localized, such that efficient decay requires an optimal distance between the terminating ribosome and the downstream EJC. When PTCs fall too close to the normal stop codon, competition from canonical termination likely suppresses NMD activation, restricting robust surveillance to PTCs positioned within a narrow optimal window (the NMD peak) of the last EJC. This quantitative PTC-to-junction distance relationship is recapitulated in transcriptome-wide data from TOPMed (Fig. 6E), suggesting it is a conserved feature across mRNAs. This quantitative PTC-to-EJC distance relationship may also underlie weak NMD observed in long exons, where a larger fraction of the exon lies beyond the ideal distance of the local EJC for optimal NMD activity.

Between the start-proximal and 3′ boundary zones, NMD operates with remarkably high and stable efficiency across internal exons of the LMNA mRNA. Within this NMD-active region (codons 25–635), the mean NMD efficiency is 0.81 ± 0.13, indicating that the vast majority of PTC-containing transcripts in this zone are degraded by the surveillance pathway. Importantly, the distribution of NMD efficiency across internal exons shows no systematic 5′-to-3′ bias (Fig. 3C), suggesting that the absolute number of EJCs upstream or downstream of a PTC does not substantially influence NMD strength in the LMNA transcript. Rather, NMD efficiency appears primarily determined by the presence of at least one downstream EJC within an optimal distance window, which is a condition satisfied throughout the internal exons. Our findings contrast with single-molecule studies suggesting EJC number influences NMD (Hoek et al., 2019); this difference may reflect transcript- or system-specific effects. Nonetheless, the high baseline efficiency in this region underscores the role of NMD as an effective quality-control mechanism for the majority of nonsense mutations.

Within internal exons, deviations from robust NMD occur only at discrete codon positions associated with specific sequence contexts, particularly TGA-C and TGA-CT motifs, rather than as gradual position-dependent changes, suggesting that a dominant source of NMD variation is likely translation readthrough. This interpretation is supported by several lines of evidence. First, the TGA-CTAG motif promotes conserved regulatory readthrough at normal termination codons in mammalian genes, which regulates functional C-terminal protein extension (Loughran et al., 2014). The shorter TGA-CT motif also stimulates readthrough broadly across evolution (Stiebler et al., 2014), and we further observe that T at the +5 position strengthens NMD attenuation. Second, eukaryotic transcriptomes selectively avoid +4 C at normal stops, consistent with evolutionary pressure to minimize basal readthrough (Cridge et al., 2006). Readthrough at PTCs in the TGA-C context in the *VHL* gene have also been reported (Buckley et al., 2024). Finally, readthrough promoting small molecules preferentially target TGA over TAG and TAA (Floquet et al., 2012; Toledano et al., 2024; Trzaska et al., 2020; Welch et al., 2007), with TGA-C serving as an especially favorable pharmacological context (Pranke et al., 2023; Toledano et al., 2024). Together, these observations support a model in which readthrough may occur at TGA-C/TGA-CT motifs at different codon positions along the transcript before NMD engagement, thereby suppressing mRNA surveillance and decay. This provides a mechanistic rationale for why the clinical efficacy of nonsense-suppression therapies varies across mutations (McDonald et al., 2017): PTCs with higher basal readthrough propensity are intrinsically more responsive to pharmacologic stimulation. In this framework, readthrough-promoting small molecules may enhance pre-existing endogenous readthrough events, selectively rescuing those PTCs already predisposed to escape NMD.

The observation that NMD variability strongly depends on PTC identity and at specific codon positions with a major influence of the residue at the +4 position (Fig. 5) strongly suggests that recognition of the stop codon by the eukaryotic release factor eRF1 during termination is a critical modulator of NMD activity. Structural studies of the eukaryotic translation termination complexes have revealed that eRF1 directly contacts both the stop codon triplet and the +4 nucleotide within the ribosomal A site (Brown et al., 2015). Recognition of the TGA context by eRF1 might be intrinsically less favorable resulting in a prolonged intermediate step before transitioning into a termination competent state. We propose that this extended dwell time has position-dependent consequences for NMD. At start-proximal PTCs, where PABPC1-mediated termination enhancement is active, slower eRF1 recognition of TGA may allow UPF1 recruitment before PABPC1-eRF3 interaction can suppress NMD, resulting in residual NMD activity. PABPC1 could additionally inhibit NMD through an interaction with eukaryotic initiation factor 4G (eIF4G), which mediates the circularization of mRNAs (Fatscher et al., 2014). In the NMD-active region, where PABPC1 influence is absent, the same slow termination kinetics instead favor readthrough, allowing ribosomes to bypass the PTC and complete translation.

Besides the position and sequence context influencing steady state abundance of PTC-containing transcripts, metabolic labeling experiments directly measuring transcriptional activity and splicing efficiency reveal that many PTCs reduce mRNA abundance through NMD-independent mechanisms, including transcription and splicing in exon 11 of the LMNA transcript (Fig. 2C). Interestingly, SMG1i treatment revealed higher expression of many variants containing PTCs in exon 1, 4, and 7 (Fig. 3A), potentially reflecting nonsense-induced transcriptional adaptation for specific PTC *LMNA* variants depending on sequence contexts.

This scenario is consistent with a prior study identifying the mouse orthologue, *Lmna*, as a hit in a screen for targets of nonsense-induced transcriptional adaptation (Mellis et al., 2024). Thus, relying solely on steady-state mRNA levels can substantially misrepresent true NMD efficiency. A clear illustration is provided by start-proximal PTCs, which showed reduced RNA abundance (Fig. 3A) despite exhibiting no measurable NMD activity (Fig. 3C). Together, these results highlight a major limitation of using only RNA-seq data under basal conditions to infer NMD rules, emphasizing the need for direct measurements to disentangle NMD from other post-transcriptional regulatory effects.

Our findings have direct implications for clinical interpretation of nonsense variants in both *LMNA* and other genes. We show that start proximal PTCs, while not triggering NMD, may also not produce any protein unless there is an optimal reinitiation context within a certain window of the PTC. This is an important factor to consider when interpreting nonsense variants in the first

200 nt of ORF, which are sometimes considered benign. Further, variants in internal exons appear to be efficiently degraded unless they occur in TGA-C or TGA-CT contexts, suggesting that such contexts may be particularly responsive to readthrough-promoting compounds, as these PTCs already exhibit reduced NMD and higher basal readthrough propensity. More broadly, our finding that the length of the start-proximal NMD inactive zone varies across genes and that dramatic NMD variations can occur at discrete codon positions underscores the need for gene-specific and sequence context characterization when interpreting nonsense variants.

## ACKNOWLEDGEMENTS

We thank Dr. David Bentley for the gift of the HAP1 parental cell line. We thank Dr. Matthew Taylor and Dr. Luisa Mestroni for helpful discussions on LMNA biology. We thank Amy Campbell, Nicole Hansen, Srinivas Ramachandran, and Ruben Rosas for insightful manuscript feedback. This work was supported by the AHA Award #1023127 (M.A.C.), TOPMed NHLBI fellowship (Z.C.A), Simons Foundation pilot award AGT011737 (Z.C.A and S.J.) University of Colorado School of Medicine Translational Research Scholars Program (S.J.), and the National Institutes of Health grant R35GM133433 (S.J.).

## AUTHOR CONTRIBUTIONS

M.A.C and S.J. conceptualized this study. M.A.C. performed all experiments, data collection, and analyses. J.S., I.E., and Z.C.A. provided curated TOPMed data used in the analyses for Figure 6. M.A.C. and S.J. performed data visualization. M.A.C. and S.J. wrote the original draft of the manuscript. All authors reviewed and approved the manuscript.

## DECLARATION OF GENERATIVE AI AND AI-ASSISTED TECHNOLOGIES

During the preparation of this work the authors used Claude.ai to improve language and readability. After using this tool, the authors reviewed and edited the content as needed and take full responsibility for the content of the publication.

## SUPPLEMENTAL INFORMATION

**Table S1.** HDR template sequences with PTC and SNV variants.

**Table S2.** List of all Cas9 gRNA sequences and primer sequences used in the study.

**Table S3.** Reference files for gDNA and cDNA read mapping.

**Table S4.** Variant mRNA expression levels before normalization and after SNV-based normalization, shown as mean ± SD of three biological replicates.

**Table S5.** Mean splicing efficiency and transcriptional activity values for all variants reported as mean ± SD of three biological replicates.

**Table S6.** Calculated NMD efficiency values before normalization and after SNV-based normalization for all variants reported as mean ± SD of three biological replicates.

## RESOURCE AVAILABILITY

### Lead contact

Further information and requests for resources and reagents should be directed to and will be fulfilled by the lead contact, Sujatha Jagannathan.

### Materials availability

All unique reagents generated in this study are available from the lead contact with a completed Material Transfer Agreement.

### Data and code availability

- All sequencing data have been deposited at GEO (xxx) and will be publicly available as of publication.
- All original code is available at the GitHub repository: https://github.com/jagannathan-lab/2025-cortazar_et_al
- Any additional information required to reanalyze the data reported in this paper is available from the lead contact upon request.

## METHOD DETAILS

### Homology-directed DNA repair (HDR) library design and cloning

SGE experiments were designed as previously described (Findlay et al., 2014). Array-synthesized DNA oligonucleotides (150 nt length) were designed for multiple SGE regions to target the full length of the *LMNA* gene. The reference LMNA gene sequence was obtained from the human genome (hg38). To enable the use of alternative CRISPR/Cas9 target sites, three synonymous substitutions were designed at three protospacer adjacent motifs (PAM) to prevent Cas9 recutting following HDR. These mutations were made at non-conserved codons to reduce the probability of perturbing functional LMNA elements. These substitutions were included in template sequences as the background in which PTC and SNV variants were introduced (Supplementary Table 1). A sequence was created for every codon substitution into each of the three stop codon types (TAA, TAG, TGA) leaving approximately ∼25 nt of unmodified sequence at the 5’ and 3’ end as adapters for cloning. Approximately 33 codons were targeted in each SGE region. For example, for a SGE region targeting 33 codons, 99 sequences were printed plus the control synonymous context without PTC (a total of 100 contexts), plus the SNV variants. The synonymous sequence context alone served as the control context for comparison. The fixed synonymous mutations additionally served to distinguish true PTCs and SNVs introduced via HDR from PCR or sequencing errors. All designed oligonucleotide contexts were synthesized by Twist Bioscience and resuspended as a single pool. Specific primers were designed to PCR amplify sequence contexts for each specific SGE target region.

For each exon, a LMNA homology plasmid was generated by PCR amplification of HAP1 gDNA (∼2 kb amplicon; 0.8–1 kb upstream and 0.8–1 kb downstream of the target exon) and cloning of products into linearized pUC19 vector using In-Fusion reactions (Takara Bioscience). LMNA homology plasmids were subsequently linearized by inverse PCR using primers with 17–25 nucleotides of overlap with amplified oligo pools specific for each SGE target region. To generate donor HDR plasmids, amplified oligo pools (30 ng) were cloned into PCR-linearized LMNA homology vectors (50 ng) using In-Fusion (Takara Bioscience). All PCR amplifications were performed with KAPA HiFi DNA Polymerase (Roche). Purification of all PCR products was performed with AMPure XP Beads (Beckman Coulter). *Escherichia coli*, Stellar competent cells (Takara) were transformed with all the In-Fusion reaction product and grown on multiple carbenicillin plates with ampicillin for 16–18 hours. All bacterial colonies were resuspended from the plates using standard Luria Broth and, after pelleting cells by centrifugation, plasmid DNA was isolated using the NucleoBond Xtra Maxiprep kit (Takara) to produce each final HDR plasmid library. All plasmid libraries were verified by Sanger and Illumina sequencing.

### CRISPR gRNA design and cloning

Protospacer CRISPR S. pyogenes Cas9 target sites were designed by choosing a non-conserved genomic site permissive to synonymous substitution within the guanine dinucleotide of the PAM or the protospacer, as well as having minimal predicted off-target activity and maximal predicted on-target activity (Doench et al., 2016; Hsu et al., 2013). Protospacer sequences were cloned into pX459 (Ran et al., 2013) (Supplementary Table 2). This plasmid expresses the gRNA from a U6 promoter, as well as a Cas9-2A-puromycin resistance (-puroR) cassette.

Complementary oligonucleotides ordered from Integrated DNA Technologies (IDT) were annealed, phosphorylated, diluted, and ligated into BbsI-digested and gel-purified pX459, as described previously (Ran et al., 2013). Ligation reactions were transformed into *Escherichia coli*, Stellar competent cells (Takara), which were plated on ampicillin-containing agar plates. Colonies were cultured and Sanger-sequenced to confirm correct gRNA sequences. Purification of sequence-verified plasmids for transfection was performed with the NucleoBond Xtra Maxiprep kit (Takara).

### Cell culture

Parental HAP1 cells were a kind gift of the laboratory of Dr. David Bentley (University of Colorado) and HAP1 LIG4 KOp cells were previously generated (Cortazar & Jagannathan, 2025). Both parental and HAP1 LIG4 KOp cell lines were cultured in medium comprising Iscove’s Modified Dulbecco’s Medium (IMDM) with L-glutamine (Gibco, catalog #12440053) and supplemented with 10% EqualFETAL Bovine Serum (Atlas Biologicals), and 1% (100 U/mL) Penicillin-Streptomycin (10,000 U/mL) (Gibco). Cells were grown in culture dishes at 37 °C with 5% CO_2_, and passaged before becoming confluent. For routine passaging, cells were washed once with 1X DPBS (Gibco), dissociated with 0.5 mL of TrypLE™ Express Enzyme (1X) (Gibco), resuspended in medium, and plated onto new plates. Cells were thawed six to seven days before transfection experiments.

### Transfection of Parent HAP1 cells

HAP1 cells were transfected using TurboFectin 8.0 (Origene) according to the manufacturer’s protocol. A 2.5X volume of Turbofectin was added to the transfection mix for each μg of plasmid DNA in Opti-MEM (Gibco). For each SGE transfection, 10 million cells were passaged to a 10-cm dish. The next day (day 0), cells were co-transfected with two pX459 plasmids encoding different sgRNAs to increase the probability of at least one inducing the DNA break within the SGE target region (3.2 μg each), plus 1.6 μg of the corresponding HDR plasmid library (8 μg total plasmid). All negative control transfections were performed for each library using a different pX459 plasmids targeting *PPP1R12C* instead of *LMNA*, thus preventing genomic integration of the library. Only for the experiment comparing the frequency of HDR events in the parent cell line versus the HAP1 LIG4 KO (Figure 1C vs. Figure S1A) an empty pX459 plasmid was used for the negative control in both cell lines. The empty plasmid approach was selected for this comparison to prevent cell death of successfully transfected cells in the NHEJ-deficient HAP LIG4 KOp cell line, since the goal was to measure any potential interference of the transfected plasmid library.

On day 1, cells were passaged into a 15 cm plate supplemented with puromycin (2 μg/mL) to select for successfully transfected cells. On day 4, cells were washed twice with 1X DPBS (Gibco) and fresh medium was added. Cell populations were maintained and passaged until sampled on day 12. Negative control transfections were harvested on the same day and were used to confirm by Illumina sequencing that PCR amplicons were not derived from the transfected plasmid DNA library. Three independent replicates were performed for each SGE experiment with a single negative control.

### Cell treatments and metabolic labelling of nascent transcripts

12 days post-transfection, SGE-treated cells were treated with 1mM SMG1i (hSMG-1 inhibitor 11j) (MedChemExpress) for 4 hours or with cycloheximide (50 μg/mL) (Sigma-Aldrich) for 6 hours, or with an equivalent volume of DMSO (Sigma-Aldrich) as vehicle control before extracting gDNA and total RNA with the AllPrep DNA/RNA kit (Qiagen).

For metabolic labeling of nascent transcripts in SGE experiment 9 (exon 11), a fraction of the cell population was incubated with 2mM 5-bromouridine (Sigma-Aldrich) for 30 minutes. At the end of the incubation time, the cell medium was removed, and cells were washed once with 1X DPBS (Gibco). Cells were resuspended in Trizol reagent (Invitrogen) and total RNA was extracted.

### Br-RNA IP and RT-qPCR targeting GAPDH nascent transcript

To validate the protocol of nascent transcript isolation, cells were incubated with 2 mM 5-bromouridine (Sigma-Aldrich) for 30 and 60 minutes. At the end of the incubation time, the cell medium was removed, and cells were washed once with 1X DPBS (Gibco). Cells were resuspended in Trizol reagent (Invitrogen) and total RNA was extracted.

Immunoprecipitation of Br-labelled nascent transcripts was performed as previously described, with minor modifications (Paulsen et al., 2014). Briefly, 50 µg of total RNA was incubated for 1 h with 2 µg of mouse anti-BrdU antibody (clone 3D4, RUO; cat. no. 555627, BD Pharmingen™) conjugated to Protein G magnetic beads (Pierce™, Thermo Scientific) in IP buffer consisting of RNase-free 1X PBS (Invitrogen) and 0.05% Triton X-100, supplemented with SUPERase·In™ RNase inhibitor (Invitrogen) and 1 mM DTT (RPI). Beads were washed four times with wash buffer (RNase-free 1X PBS containing 0.05% Triton X-100). Enriched nascent transcripts were eluted by heating at 90 °C for 5 min. RNA concentration was quantified using the Qubit™ RNA High Sensitivity assay (Invitrogen).

For analysis of GAPDH nascent transcripts, cDNA was synthesized from 40 ng of enriched Br-labelled RNA using SuperScript™ II reverse transcriptase (Invitrogen) and N6 random primers. RT–qPCR was performed using genomic DNA to generate standard curves for primers targeting intergenic, exonic, and intronic regions, whereas cDNA derived from total RNA was used to generate a standard curve for primers spanning GAPDH exon–exon junctions. Primer sequences are provided in Supplementary Table 2.

### In-vitro synthesis and labeling of spike-in RNA

The spike-in RNA was transcribed by T7 RNA Polymerase (Invitrogen) from a gBlock (IDT) DNA template following the manufacturer’s protocol. Briefly, synthesis was performed in 1X T7 transcription buffer with 0.5mM ATP, GTP and CTP, 0.05 mM UTP, 0.2 mM Br-UTP, 10mM DTT and 2 ng/μL template, supplemented with RNaseOUT™ Recombinant Ribonuclease Inhibitor (Invitrogen) in nuclease-free water at 37°C for 60 minutes. The sequence of the spike-in can be found in the primer list (Supplementary Table 2).

### Multiplex RT-PCR gel electrophoresis

Nascent transcripts were labelled with bromouridine as described above in wild-type HAP1 cells. For immunoprecipitation of nascent transcripts, 10 μg of input total RNA was spiked with increasing amounts of *in vitro*-synthesized spike-in RNA (0, 0.1, 1, 10 pg) or 10 pg of unlabelled spike-in. Immunoprecipitation was performed as described above but using 15 μL of Protein G magnetic beads (Pierce Thermo Scientific) and 1 μg of Mouse Anti-BrdU (Clone 3D4, RUO; Cat 555627, BD Pharmingen™).

To amplify unspliced (exon 11-intron 11) and spliced LMNA (exon 11-exon 12) from nascent transcripts including the spike-in RNA, a multiplex cDNA synthesis followed by multiplex PCR approach was performed. For each sample, two primers were used in the same reaction for cDNA synthesis to prime at both exon 12 (5’-GGGCAGGGGGTGGGCATG-3’) and at intron 11 (5’-GCTCAGACAAGAGGGGCAGG-3’) using 20 ng of nascent transcripts in a 20 μL reaction with SuperScript™ II Reverse Transcriptase (Invitrogen). The spike-in is primed by the primer recognizing the intronic sequence for cDNA synthesis of the spike-in.

A multiplex PCR reaction using the cDNA product as template was performed to amplify the unspliced, spliced, and spike-in products. A common forward primer binding on exon 11 (5’-CAGGAGCCCAGGTGGGCG-3’) and two different reverse primers binding exon 12 (5’-AGGTGAGGAGGACGCAGGAAG-3’) or intron 11 (5’-GCAGAGGTGGGCTGTCTAGGAC-3’) were used. The forward and the reverse primer binding to the intronic sequence amplify the spike-in signal. PCR products were then run on an agarose gel and visualized.

All primers were first optimized with nascent transcripts purified from wild-type HAP1 cells. Individual cDNA primers were validated independently and in combination in the multiplex approach by gel electrophoresis and Sanger sequencing. Similarly, primer pairs for PCR were validated independently and in combination in the multiplex approach by gel electrophoresis and sequencing. All PCR reactions were performed with KAPA HiFi DNA polymerase (Roche).

### Multiplex library preparation from nascent transcripts

To create sequencing libraries from SGE-treated cells (SGE experiment 9, exon 11), nascent transcripts were immunoprecipitated from 50 μg of RNA as described above (Br-RNA IP RT-qPCR), and the same multiplex cDNA and multiplex PCR procedures described above, were performed. cDNA synthesis was performed with 100 ng of enriched nascent transcripts in a total reaction volume of 20 μL. The entire cDNA product was amplified in multiple PCR reactions using 2.5 μL as template in 50 μL PCR reactions. PCR reactions from the same sample were pooled, mixed thoroughly and DNA fragments were purified with AMPure XP Beads (Beckman Coulter).

As for gDNA and cDNA libraries, two additional PCR reactions (second and third PCRs) were performed. The second PCR corresponds to a nested PCR to add universal Illumina sequencing adapters (P5, P7) common to unspliced and spliced amplicons using the PCR products from the first multiplex PCR. A common forward primer binding exon 11 was used (5’-ACACTCTTTCCCTACACGACGCTCTTCCGATCTAGTGTCACGGTCACTCGCAG-3’) and two different reverse primers binding at exon 12 (5’-AGACGTGTGCTCTTCCGATCTGGTCCCAGATTACATGATGCTGCAG-3’) and at intron 11 (5’-AGACGTGTGCTCTTCCGATCTCCTGCAGGATTTGGAGACAAAGC-3’). At this step, all amplicons, unspliced and spliced are of 150 nucleotides in length and only differ at the 3’ end (5’ splice site) with intronic or exon sequence, plus the adapter sequence. Finally, unique dual indexes were added by PCR (third PCR) using primers with 5’ extensions binding at the adapter sequence. All sequencing libraries were purified with AMPure XP Beads (Beckman Coulter) and concentrations were determined with the Qubit dsDNA High Sensitivity kit (Invitrogen).

### Nucleic acid sampling and sequencing library production

At day 12 post-transfection, DNA and total RNA were purified at the end of treatments (i.e. DMSO, SMG1i, cycloheximide) from the same cell population using the AllPrep DNA/RNA kit (Qiagen). DNA and RNA samples were quantified with the Qubit dsDNA or RNA High Sensitivity kits (Invitrogen).

PCR primers for genomic DNA were designed such that one primer would anneal outside of the homology arm sequence, thereby selecting for amplicons derived from gDNA and not plasmid DNA (first PCR) (all primer sequences are reported in Supplementary Table 2) PCR conditions (KAPA HiFi) were optimized on wild-type HAP1 gDNA and the lowest number of cycles was used to amplify the SGE target region from 150 ng of template (18–22 cycles). A total of 8 μg gDNA (or all of the extracted gDNA but no less than 6 μg) was amplified in multiple 50 μL reactions to ensure adequate sampling. After PCR, all 50 μL reactions were pooled, mixed thoroughly, and a fraction was purified using AMPure XP Beads (Beckman Coulter).

A nested PCR (second PCR) was performed using 5 ng of product from the first PCR reaction as template and PCR primers with 5’ universal extensions to add part of Illumina adapter sequences (P5, P7). The HDR plasmid libraries were also PCR-amplified at this step starting from 10 ng of plasmid DNA. PCR products were purified using AMPure XP Beads (Beckman Coulter). Finally, unique dual indexes were added by PCR (Third PCR) using primers with 5’ extensions and priming to the universal adapter sequence.

A total of 20 μg of RNA was reverse transcribed per sample using SuperScript™ II Reverse Transcriptase (Invitrogen) according to the manufacturer’s protocol. Briefly, four reactions were performed each containing 5 μg of RNA priming in exon 12 of the LMNA mRNA (oligo sequence 5’-GGGCAGGGGGTGGGCATG-3’) in 50 μL reaction. For SGE experiments targeting exon 1, priming was performed in exon 10 (oligo sequence 5’-TCCATCCTCATCCTCGTCGTCC-3’). All four reaction products were pooled and mixed. PCR Primers were designed for each exon to amplify across exon junctions (First PCR), which excludes amplification from any potential contaminating gDNA. PCR conditions were optimized on wild-type HAP1 cDNA and primer pairs yielding a single PCR product were selected. The lowest number of cycles was performed to reach amplification (20–22 cycles). The entire cDNA reaction (200 μL) was used as template for PCR amplification using 2.5 μL per 50 μL reaction. After PCR, all 50 μL reactions were pooled, mixed thoroughly, and a fraction was purified using AMPure XP Beads (Beckman Coulter).

A nested PCR was performed to amplify the SGE target region (second PCR) using 5 ng of the product from the first PCR reaction as template and PCR primers with 5’ universal extensions to add part of Illumina adapter sequences (P5, P7). PCR products were purified using AMPure XP Beads (Beckman Coulter). Finally, unique dual indexes were added by PCR (third PCR) using primers with 5’ extensions. All sequencing libraries were purified with AMPure XP Beads (Beckman Coulter) and concentrations were determined with the Qubit dsDNA High Sensitivity kit (Invitrogen).

### Sequencing and read alignment

Libraries were diluted to 10 nM, pooled in equimolar amounts, and sequenced on an Illumina NovaSeq platform using paired-end sequencing (Novogene) (150 cycles for read 1 and 150 cycles for read 2). This sequences the insert corresponding to the full 150-nt SGE-target region from both strands of the DNA fragment. Ten million reads were allocated to each gDNA and cDNA sample and five million reads for each HDR library and negative control sample. To filter out sequencing errors we used SeqPrep (https://github.com/jstjohn/SeqPrep).

Sequencing errors would cause a mismatch between the complementary read 1 and read 2, which can be filtered out with SeqPrep. The following parameters were used to merge perfectly matched overlapping read pairs for the entire 150 nt length: -m 150 -L 150 -o 150 -q 0”. The filtered merged reads were used for alignment of HDR sequence contexts (variants). To analyze indel frequencies, SeqPrep was performed with the following parameters, which do not exclude inserts not conforming to 150 nt: “-A GGTTTGGAGCGAGATTGATAAAGT-B CTGAGCTCTCTCACAGCCATTTAG, -m 20 -L 20 -o 20 -q 0”.

Merged SeqPrep filtered reads of 150 nt were aligned using Bowtie (Langmead et al., 2009). The default parameters were used, except that “-v 0” was included to retain only perfectly matching reads. Reads from each gDNA or cDNA library were aligned to a gDNA or cDNA reference correspondingly (reference files are provided in Supplementary Table 3). gDNA and cDNA references may vary given that some 150 SGE target regions targeting the edge of an exon (to include all codons at the 5’ or 3’) include intronic sequences. cDNA references for those experiments replace the intronic sequence with the corresponding exon-exon junction sequence. Reads in libraries made from nascent transcripts were aligned to a single reference containing the exon 11-intron 11 boundary (unspliced reads) and the exon 11-exon 12 junction (spliced reads) using the same parameters for alignment as above. For SGE experiments 1, 4 (exon 1), 5, 6 (exon 4), a small fraction of reads (< 15%) failed to align due to one synonymous mutation not being integrated by HDR. For these regions, we appended alternative reference sequences containing only the two synonymous mutations closest to the Cas9 break and repeated the alignment. Total counts for each variant were calculated with a Python script.

### Calculation of mRNA expression levels and variant filtering

To calculate mRNA expression levels, first the count for each variant in the cDNA library was normalized to the frequency of genomic integration of each variant and then normalized to the value of the synonymous context. Using a Python script, the count for each variant in the cDNA library was divided by the count in the gDNA. This value for each mutation was then divided by the value obtained for the synonymous context. To compare across different SGE experiments an R script was used to perform SNV-based normalization. To do this, the median expression level of all SNV variants in each experiment was used to divide the mean mRNA expression levels of each variant within the same experiment. Variant mRNA expression levels were reported before normalization (Fig. S4A) and after SNV-based normalization (Fig. 3A) and are shown as mean ± SD of three biological replicates. Calculated values are provided in Supplementary Table 4.

To filter low-confidence variants, only reads passing SeqPrep filtering were retained. Variants supported by fewer than 10 aligned reads in either gDNA or cDNA libraries were excluded. The majority of variants (>90%) were supported by more than 1,000 reads. Only variants present in both DMSO and SMG1i conditions were retained. Variants with high replicate variability (SD > 0.2) in either condition were also excluded.

### Calculation of transcriptional activity, splicing efficiency and variant filtering

To calculate transcriptional activity for exon 11 variants, unspliced and spliced counts from multiplex libraries made from nascent transcripts were summed for each variant, normalized to genomic integration frequency (gDNA library), and then normalized to the synonymous context. Mean expression levels across replicates are reported as mean ± SD of three biological replicates.

Splicing efficiency was calculated as the fraction of spliced reads over total reads and normalized to the synonymous context. Mean splicing efficiency values are reported as mean ± SD of three biological replicates. All exon 11 variants were supported by >1,000 reads and showed low variability (SD < 0.2) and were retained. Values are provided in Supplementary Table 5.

### Calculation of NMD efficiency and variant filtering

To determine NMD efficiency, the expression level of each variant relative to the synonymous context upon SMG1i treatment was calculated from cDNA libraries using a Python script.

Specifically, the counts for each variant in the DMSO and SMG1i treatment libraries were divided by the count of the synonymous context within the same library. Then, these values were used in the following formula: NMD efficiency = 1 – (DMSO/SMG1i). To compare NMD efficiency across different SGE experiments an R script was used to perform SNV-based normalization. For this, the median NMD efficiency value of all SNV variants (close to zero) in each experiment was subtracted from the NMD efficiency value for each PTC variant within the same experiment. The NMD efficiency value for each PTC variant was reported before SNV-based normalization (Fig. S4C) and after normalization (Fig. 3C) and all values are shown as mean ± SD of three biological replicates.

To filter low confidence variants only reads that passed SeqPrep-based sequencing error filtering were kept for alignment. A small minority of sequence contexts supported by fewer than 10 aligned reads either in the DMSO or SMG1i cDNA libraries were discarded from the analysis. The vast majority of variants (>90%) were supported by more than 1,000 reads in cDNA libraries. Only variants present in both DMSO and SMG1i treatment conditions were kept for further analysis. Variants for which the calculation of mean expression displayed substantial replicate variation (SD > 0.2) in either DMSO or SMG1i treatment libraries were also excluded from the analysis. Calculated NMD efficiency values can be found in Supplementary Table 6.

### TOPMed Allele-specific expression analysis

RNA TOPMed cis-e/sQTL results were generated in a collaboration between the TOPMed Informatics Research Center, TOPMed Multi-Omics working group, and the TOPMed parent studies contributing RNA-seq and distributed to TOPMed investigators. For the accurate calculation of allele specific expression (ASE) from RNA-Seq data, we used STAR (Dobin et al., 2013) aligner’s WASP functionality (van de Geijn et al., 2015) to mitigate the reference bias while mapping the RNA-Seq reads to the reference genome. Then we used GATK ASEReadCounter version (v3) (https://gatk.broadinstitute.org/hc/en-us/articles/360037428291-ASEReadCounter)(Castel et al., 2020) to extract allele specific expression for truncating variants. NMD efficiency was then calculated as the ratio of reference read allele count to total allele count, sampled 100 times across individuals with a particular variant, and the median ASE value used for plotting (Rivas, Pirinen, Conrad, Lek, Tsang, Karczewski, Maller, Kukurba, DeLuca, Fromer, Ferreira, Smith, Zhang, Zhao, Banks, Poplin, Ruderfer, Purcell, Tukiainen, Minikel, Stenson, Cooper, Huang, Sullivan, Nedzel, Bustamante, et al., 2015). Variant consequences and transcript mappings were annotated on GRCh38 using Ensembl VEP v106 with dbNSFP (Liu et al., 2020) and plugin suite.

## QUANTIFICATION AND STATISTICAL ANALYSIS

### Data analysis, statistical tests, and visualization

All experiments were performed with a sample size of n=3. The exact code and packages used for these analyses can be found in this repository: https://github.com/jagannathan-lab/2025-cortazar_et_al

## SUPPLEMENTAL INFORMATION

### SUPPLEMENTARY FIGURES

**Supplementary figure 1.**
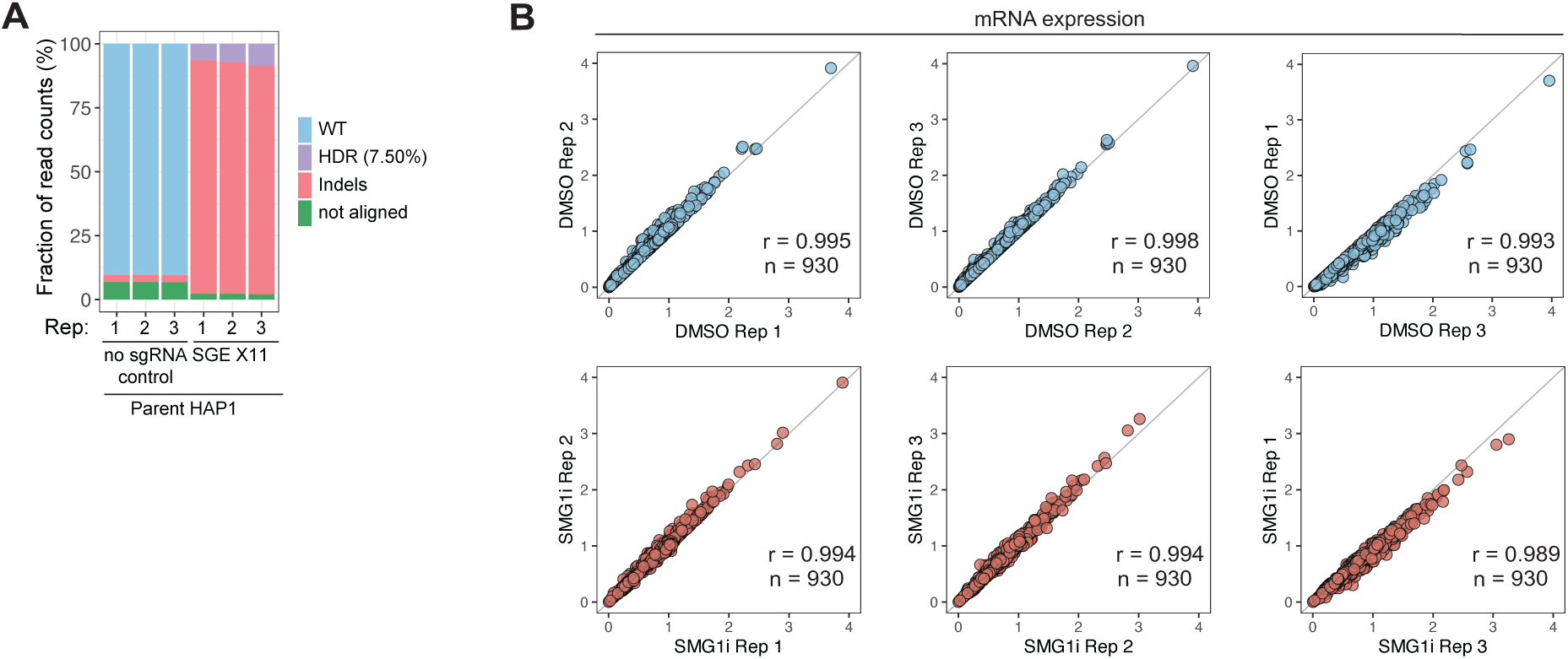
**(A)** Proportion of sequenced read tags from genomic DNA mapping to wild-type (WT; unedited cells), homology-directed repair (HDR), or insertions and deletions (indels) in parental HAP1 cells with and without sgRNA. **(B)** Correlation of mRNA expression levels between replicates in DMSO and SMG1i conditions for each stop codon type (TAA, TAG, TGA).

**Supplementary figure 2.**
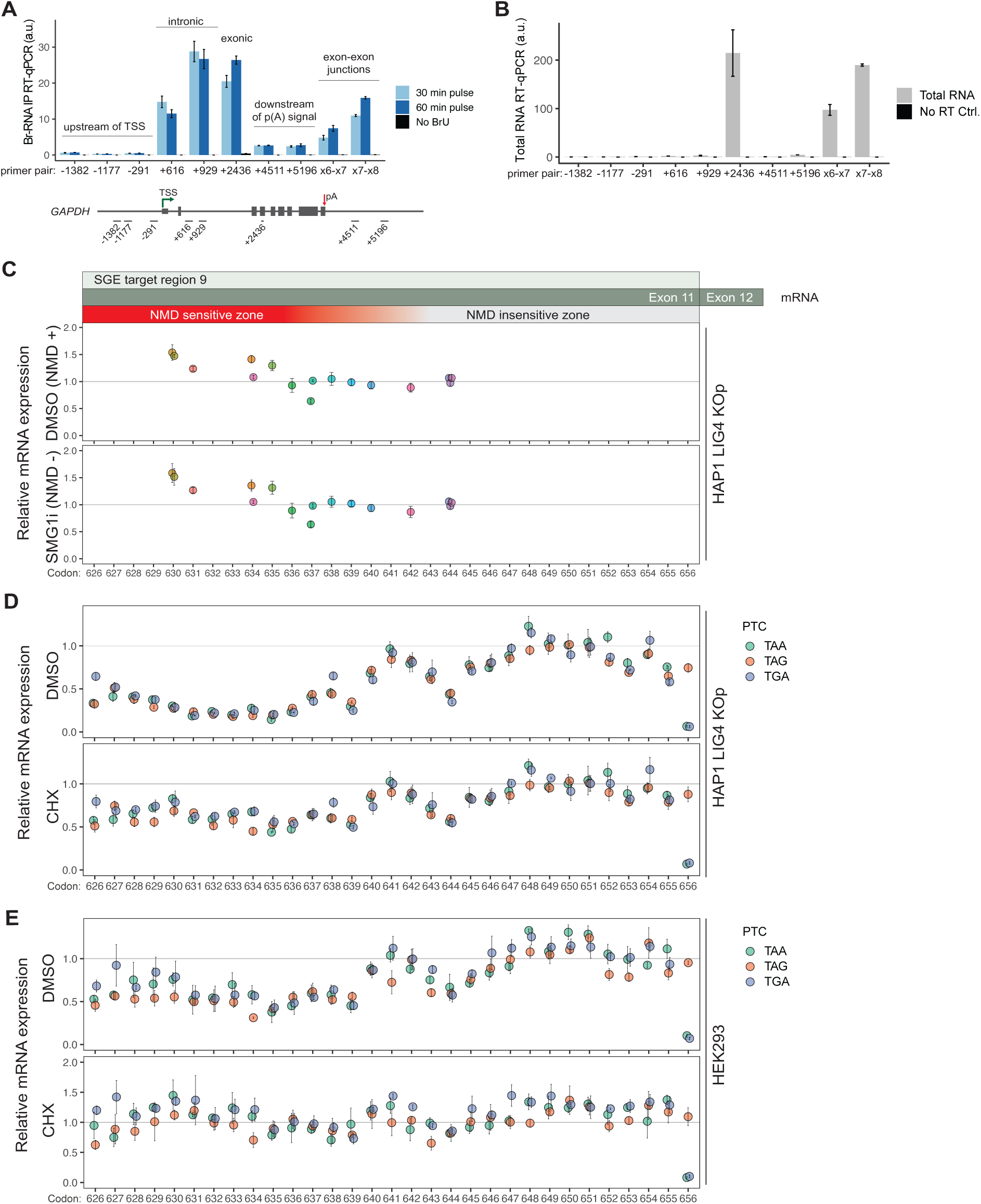
(A-B) Validation of BrU labeling and immunoprecipitation of nascent RNAs via RT-qPCR of GAPDH pre-mRNA signal from (A) nascent transcripts, or from (B) total, input RNA. (C) mRNA expression of *LMNA* variants containing SNVs at the indicated codon position, relative to the synonymous sequence control, under DMSO or SMG1i conditions. (D-E) mRNA expression of *LMNA* variants containing a single PTC at the indicated codon position, relative to the synonymous sequence control, under DMSO or cycloheximide (CHX) conditions in Hap1 (D) or HEK293 cells (E).

**Supplementary figure 3.**
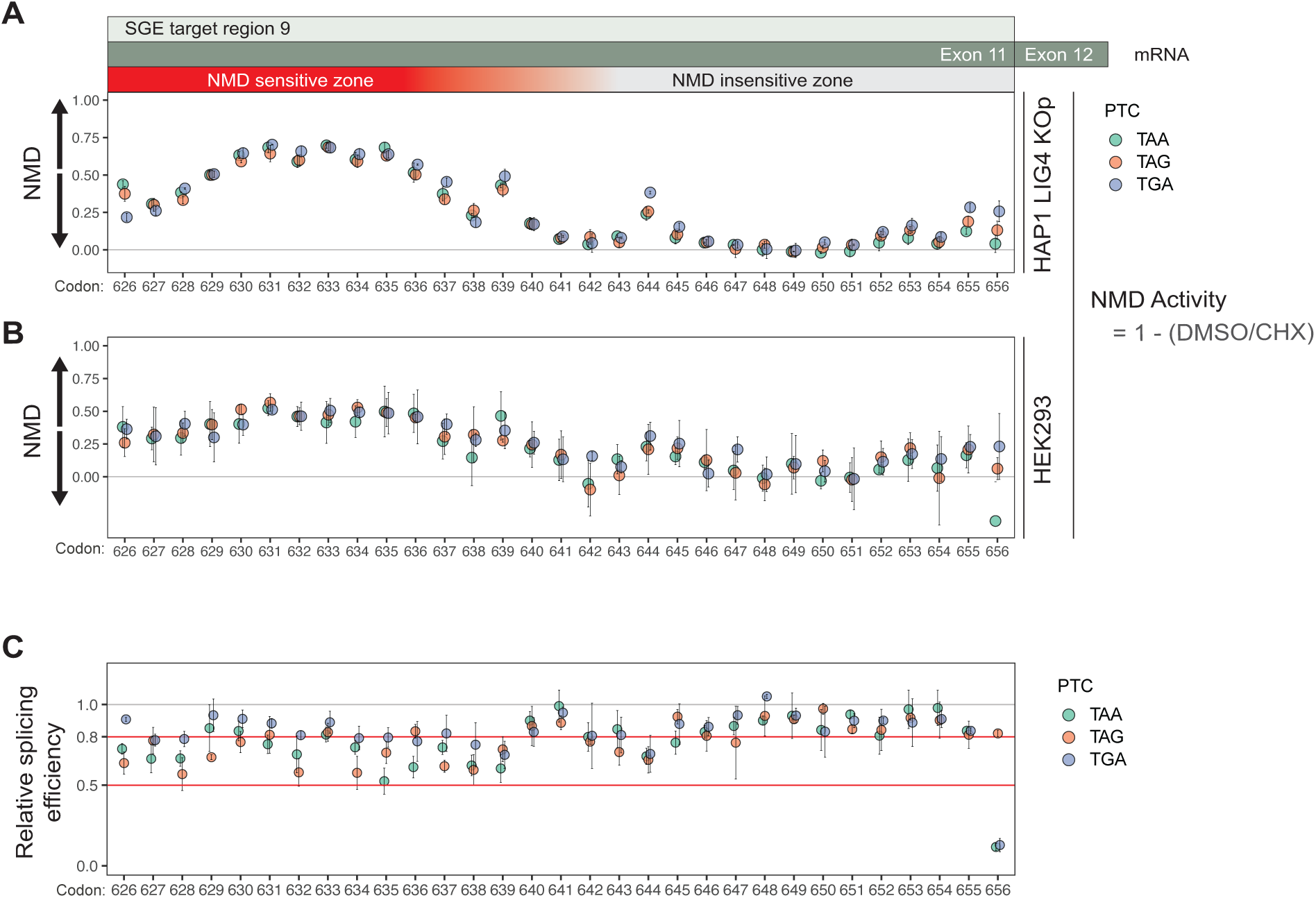
(A) NMD efficiency of *LMNA* variants containing a single PTC at the indicated codon position, under DMSO or Cycloheximide (CHX) conditions in Hap1 LIG4 knock out cell population. (B) Same as (A) in HEK293 wild-type cells. (C) Same plot as Fig 2E, with a modified Y-axis to highlight differences in relative splicing efficiency.

**Supplementary figure 4.**
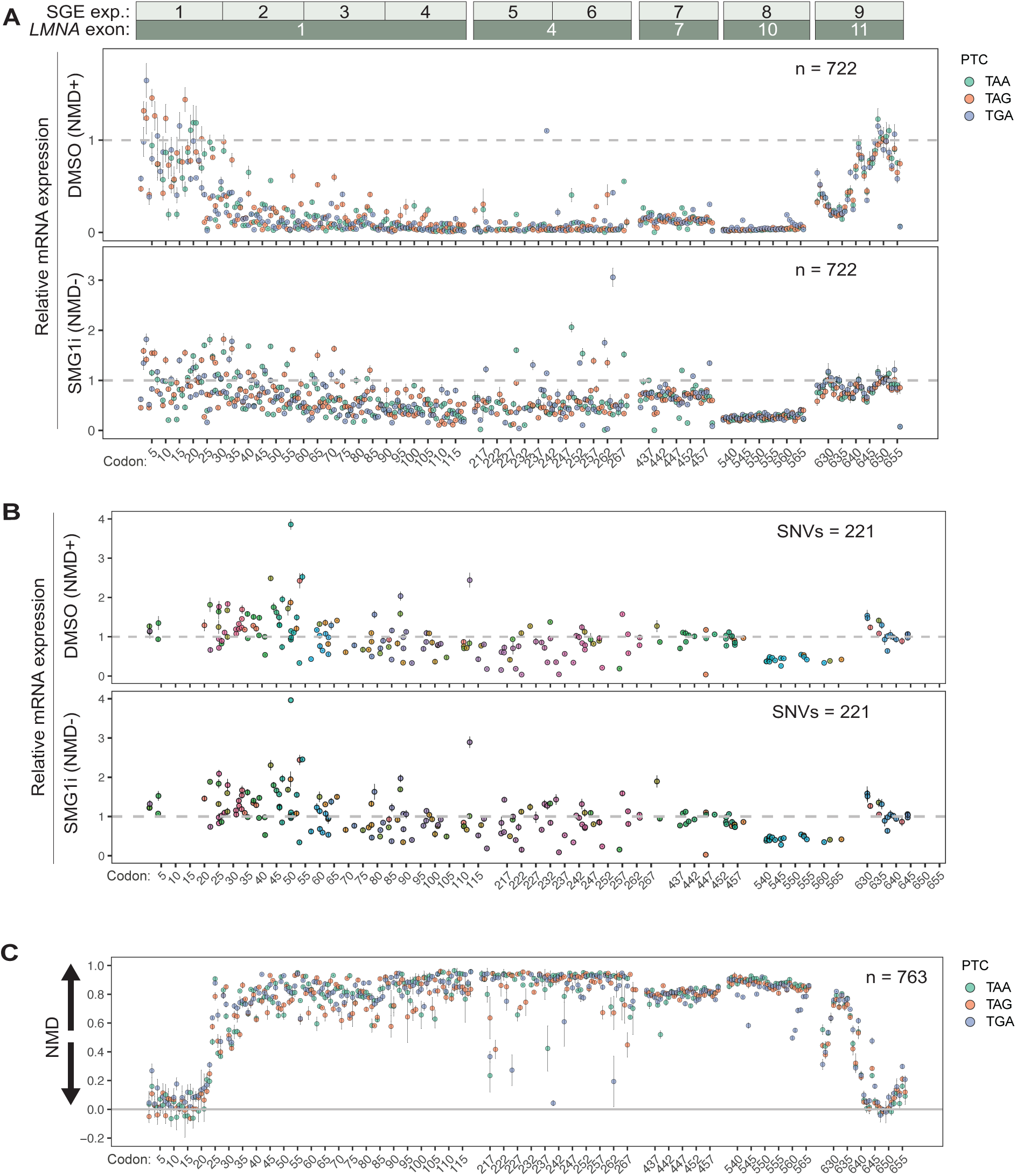
(A) mRNA expression values relative to the control, synonymous sequence context before SNV-based normalization. (B) mRNA expression levels of SNV variants relative to the synonymous sequence context. (C) NMD efficiency before SNV-based normalization.

**Supplementary figure 5.**
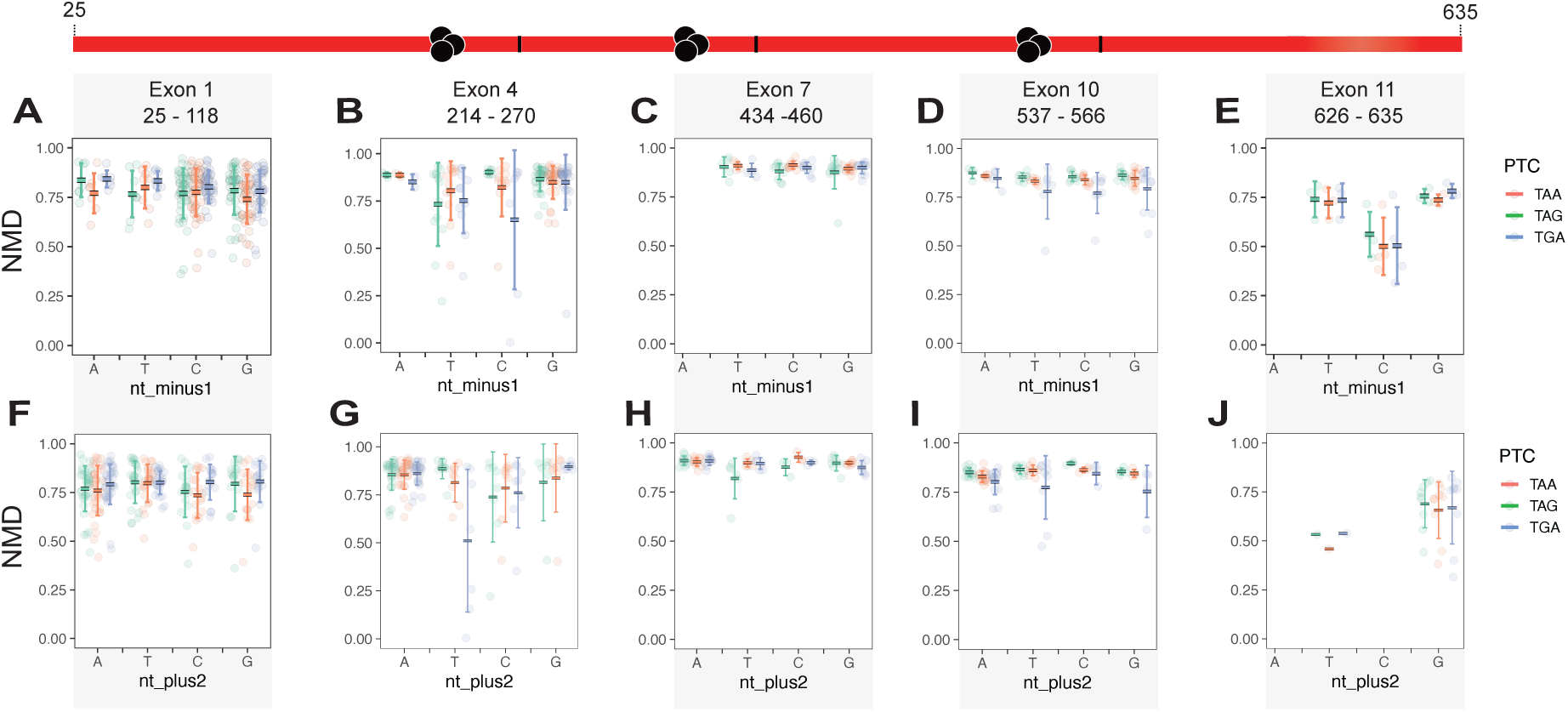
Mean NMD efficiency as a function of the nucleotide identity at positions −1 (A-E) and +5 (F-J) relative to the PTC within the NMD-active region, analyzed separately for each exon in the NMD-active region.

